# Elav-mediated exon skipping and alternative polyadenylation of the *Dscam1* gene is required for axon outgrowth

**DOI:** 10.1101/613059

**Authors:** Z. Zhang, K. So, R. Peterson, M. Bauer, H. Ng, Y. Zhang, J.H. Kim, T. Kidd, P. Miura

## Abstract

Many metazoan genes express alternative long 3′ UTR isoforms in the nervous system, but their functions remain largely unclear. In *Drosophila melanogaster*, the *Dscam1* gene generates short and long (*Dscam1-L*) 3′ UTR isoforms due to alternative polyadenylation (APA). Here, we found that the RNA-binding protein Embryonic Lethal Abnormal Visual System (Elav) impacts *Dscam1* biogenesis at two levels, including regulation of long 3′ UTR biogenesis and skipping of an upstream exon (exon 19). MinION long-read sequencing confirmed the connectivity of this alternative splicing event to the long 3′ UTR. Knockdown or CRISPR deletion of *Dscam1-L* impaired axon growth in *Drosophila*. The *Dscam1* long 3′ UTR was found to be required for correct Elav-mediated skipping of exon 19. Elav thus co-regulates APA and alternative splicing to generate specific *Dscam1* transcripts that are essential for neural development. This coupling of APA to alternative splicing might represent a new class of regulated RNA processing.

**Graphical Abstract:** 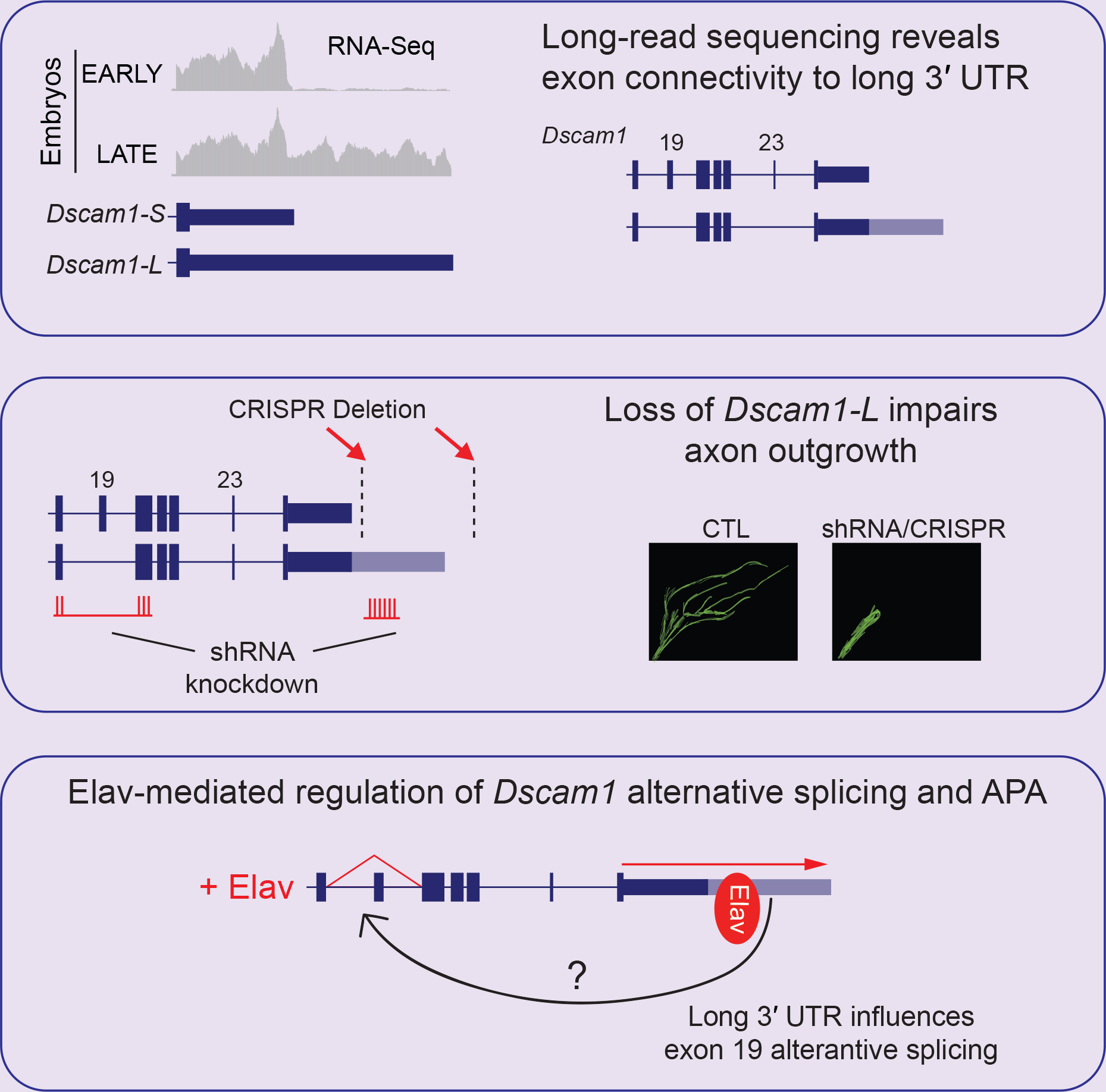

**Highlights:**

- Elav regulates *Dscam1* long 3′ UTR (*Dscam1-L*) biogenesis
- Long-read sequencing reveals connectivity of long 3′ UTR to skipping of upstream exon 19
- Loss of *Dscam1-L* impairs axon outgrowth
- *Dscam1* long 3′ UTR is required for correct splicing of exon 19

## Introduction

Alternative polyadenylation (APA) is a key event in RNA processing that most commonly results in mRNAs with different length 3′ UTRs (tandem APA or 3′ UTR APA) (Miura et al., 2014; Tian and Manley, 2017). Well over half of genes in *Drosophila*, zebrafish, mice and humans undergo APA (Hoque et al., 2013; Lianoglou et al., 2013; Sanfilippo et al., 2017; Smibert et al., 2012; Ulitsky et al., 2012). APA generates alternative length 3′ UTRs depending on tissue and cell type, with testis generating short 3′ UTR isoforms, and brain tissue generating extended or long 3′ UTR isoforms (Miura et al., 2013; Ramskold et al., 2009; Sanfilippo et al., 2017; Smibert et al., 2012).

Previous RNA-Seq studies have uncovered that 3′ UTR extension or lengthening is a pervasive event in metazoan nervous systems. In *Drosophila*, hundreds of genes have been found to express alternative long 3′ UTR isoforms in neural tissues (Brown et al., 2014; Smibert et al., 2012). In mice and humans, thousands of genes were found to express previously unannotated long 3′ UTR isoforms (Miura et al., 2013). Long 3′ UTR isoforms tend to be associated with lower molecular weight polysomal fractions than their shorter counterparts, suggesting they are less efficiently translated (Blair et al., 2017). Elements within 3′ UTRs are also important for localization to dendrites and axons (Cioni et al., 2018; Glock et al., 2017).

Alternative long 3′ UTRs harbor increased real estate compared to their short counterparts for regulation by RNA-binding proteins (RBPs) and microRNAs that can control mRNA stability, localization and translation (Miura et al., 2014). Several long 3′ UTR isoforms have been previously implicated in neural development. An alternative long 3′ UTR isoform of *Impa1* directs mRNA localization to axons, and its knockdown by siRNAs leads to axonal degeneration in rat sympathetic neurons (Andreassi et al., 2010). Other studies have used indirect methods to abrogate alternative 3′ UTR isoforms. For instance, to study the function of long *BDNF* transcripts in mice, a SV40 polyadenylation (polyA) site was inserted downstream of the BDNF proximal polyadenylation site to inhibit expression of the long 3′ UTR (An et al., 2008). These mice had impaired synaptic transmission and exhibited hyperphagic obesity (An et al., 2008; Liao et al., 2012). More recently, the function of the long 3′ UTR isoform of CamKII was studied indirectly by generating a CamKII knockout that continued to express short 3′ UTR CamKII via maternal contribution (Kuklin et al., 2017). These flies displayed impaired synaptic plasticity, which was at least partially attributed to impaired local translation of CamKII.

The neuronal RBP embryonic lethal abnormal visual system (Elav) binds to U-rich elements to regulate alternative splicing and APA (Soller and White, 2003; Zaharieva et al., 2015). Elav has been proposed to compete with the cleavage and polyadenylation machinery for the downstream U-rich element (DUE) found at proximal polyadenylation sites, thus promoting long 3′ UTR biogenesis (Hilgers et al., 2012). This mechanism has been described for other RBPs in regulating alternative polyadenylation (Gawande et al., 2006; Mansfield and Keene, 2012; Zhu et al., 2007). In addition, a role for Elav binding to gene promoters has also been implicated in the mechanism of 3′ UTR lengthening (Oktaba et al., 2015).

The *Drosophila* Down syndrome cell adhesion molecule (*Dscam1*) gene encodes a transmembrane receptor that plays an important role in neurite self-avoidance, axon guidance, and maintenance of neural circuits (Schmucker and Chen, 2009; Zipursky et al., 2006). *Dscam1* expresses two 3′ UTR variants, a short 3′ UTR of ~1.1 kb (*Dscam1-S*), and a long 3′ UTR variant of ~2.8 kb (*Dscam1-L*) (Smibert et al., 2012). *Dscam1* is appreciated as the most extensively alternatively spliced gene known in nature, with the potential to generate over 38,000 mRNA protein isoforms (Brown et al., 2014; Schmucker et al., 2000). With advances in long-read sequencing it has become possible to identify mRNA alternative exon connectivity in an unambiguous way. MinION long-read RNA-sequencing of the three *Dscam1* hypervariable exon clusters 4, 6, and 9, which are important for dendritic self-avoidance (Hughes et al., 2007; Matthews et al., 2007), identified at least 7,874 unique splice-forms (Bolisetty et al., 2015). In addition to these clusters, alternative splicing of exons 19 and 23 generates endodomain diversity (Yu et al., 2009). Suppression of *Dscam1* mRNAs lacking exons 19 or 23 was previously found to inhibit postembryonic neuronal morphogenesis, demonstrating the crucial importance of skipping these exons for *Dscam1* function in neurons (Yu et al., 2009). Despite their importance, the factors that regulate alternative splicing of exons 19 and 23 are unknown.

In this study, we set out to determine the functional impact of long *Dscam1* 3′ UTR loss on neural development. We found that Elav promotes *Dscam1* long 3′ UTR biogenesis, which restricts its expression to neurons. We specifically knocked down *Dscam1-L* by short hairpin RNA (shRNA) in neurons, and found that this resulted in severely compromised locomotion and adult lethality. Overall Dscam1 protein levels remained unchanged in the knockdown condition. This prompted us to investigate upstream splicing events that coincide with the expression of the long 3′ UTR. We identified that *Dscam1-L* transcripts preferentially exclude exon 19. Knockdown of *Dscam1-*L severely impaired mushroom body bifurcation and suppressed axon outgrowth of small ventral lateral neurons (sLNvs). The importance of *Dscam1-L* for axon outgrowth was confirmed in flies harboring a CRISPR/Cas9-mediated deletion of the long 3′ UTR region. We found that the skipping of exon 19 is mediated by Elav, and this skipping event is deregulated upon loss of the long 3′ UTR. In summary, we have found that Elav regulates *Dscam1* at both the levels of alternative splicing and alternative polyadenylation, and the resulting transcripts that bear the long 3′ UTR and lack exon 19 are required for axon outgrowth.

## Results

### Elav regulates biogenesis of *Dscam1-L*

*Dscam1* expresses two 3′ UTR variants, a short 3′ UTR of ~1.1 kb (*Dscam1-S*), and long 3′ UTR variant of ~2.8 kb (*Dscam1-L*). We sought to determine the mechanism that regulates biogenesis of *Dscam1-L*. Visual examination of publicly available RNA-Seq tracks (See methods) using Integrated Genomics Viewer (Thorvaldsdottir et al., 2013) suggested that the *Dscam1* long 3′ UTR isoform is not expressed in early-stage embryos, but appears in late-stage embryos which coincides with the development of the nervous system. The long 3′ UTR is also expressed in larval stage 3 (L3) central nervous system (CNS) (Figure 1A). To confirm these trends, we monitored *Dscam1* 3′ UTR isoforms by Northern analysis using a probe hybridizing to the constitutively expressed *Dscam1* exon 11. We found that late stage embryos (8-12 hr, 12-16 hr after egg laying) exhibited expression of *Dscam1-L* whereas early stage embryos did not (0-4 hr, 4-8 hr) (Figure 1B). There have been reports that 3′ UTRs can be cleaved and form stable fragments separated from their upstream protein-coding regions (Kocabas et al., 2015; Malka et al., 2017). Northern blot using a probe targeting the 3′ UTR downstream of the stop codon did not reveal evidence for such isolated 3′ UTRs, although this does not preclude their existence (Figure S1). Western analysis showed that late stage embryos express enhanced levels of Elav protein compared to early stages, which coincides with the expression of *Dscam1-L* (Figure 1B).

**Figure 1:**
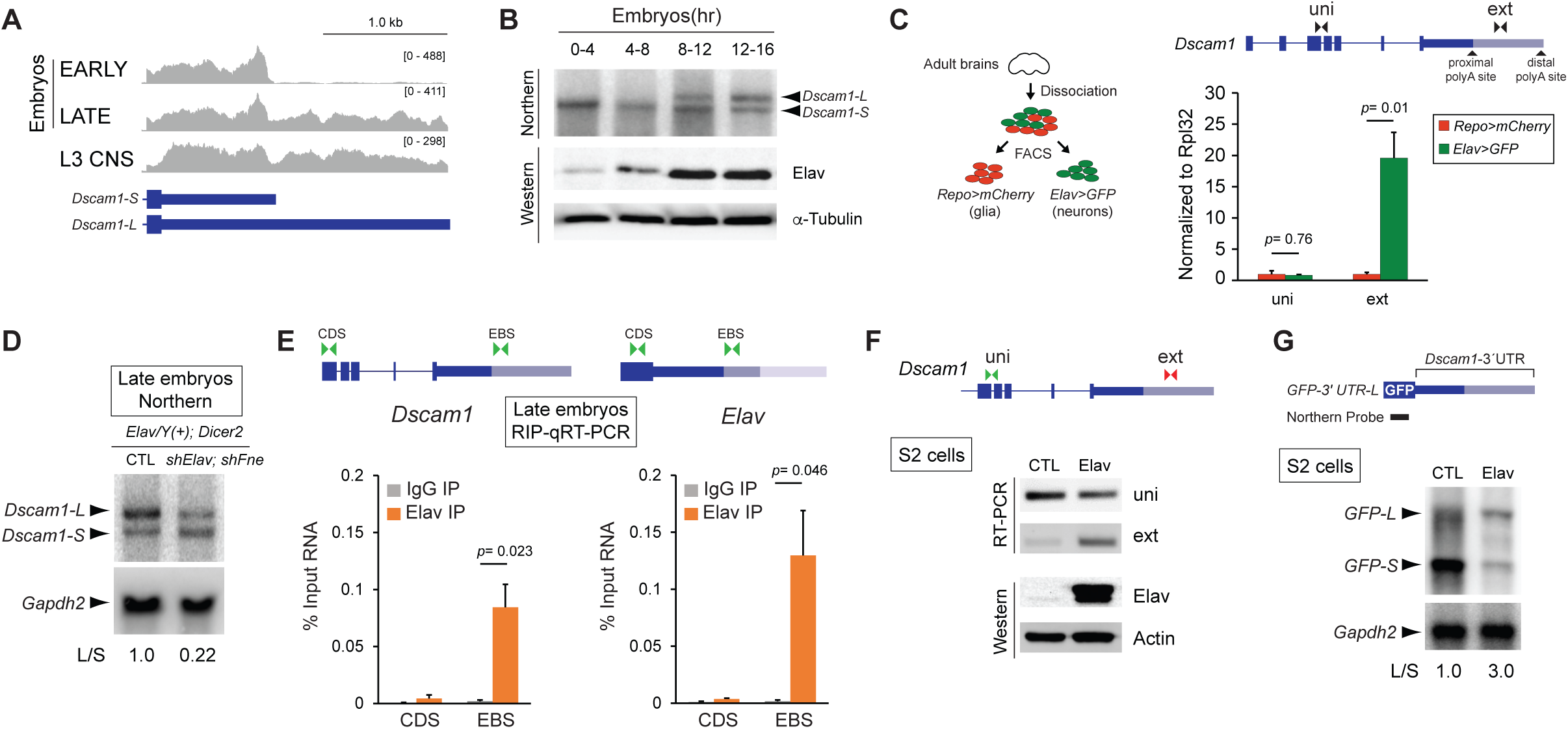
Elav regulates *Dscam1* long 3′ UTR biogenesis. **A)** RNA-Seq tracks demonstrate extension of *Dscam1* 3′ UTR is absent in EARLY (6-8 hrs after egg laying) embryos, but is induced in LATE embryos (16-18 hrs) and L3 larval CNS. **B)** Northern analysis shows expression of *Dscam1* long 3′ UTR isoform (*Dscam1-L*) in late stage embryos whereas the short 3′ UTR isoform (*Dscam1-S*) is expressed throughout development (top panel). Northern probe was designed to target common *Dscam1* exon 11. Western blot shows increased expression of Elav in the later embryonic time points. α-Tubulin is shown as a loading control. **C)** FACS analysis of adult brains from Repo positive cells (Glia) and Elav positive cells (Neurons). qRT-PCR detection of both the short and long *Dscam1* transcripts using uni primers shows similar levels between Repo and Elav positive cells. Detection using ext primers demonstrates that *Dscam1-L* is highly enriched in Elav positive cells. Error bars represent SEM (standard error of the mean). n=3. *P* value reflects unpaired Student’s T test. **D)** Northern analysis of 16-20 hr embryos shows that knockdown of Elav and the related protein FNE by shRNAs in neurons resulted in a reduction in the ratio of *Dscam1-L* to *Dscam1-S* (L/S). *Elav* represents *Elav-GAL4*, and *Dicer2* represents *UAS-Dicer2*. *Dscam1* Northern probe targets common exon 11. Gapdh2 Northern is shown as a loading control. **E)** RIP-RT-qPCR experiments demonstrate binding of Elav downstream of the *Dscam1* proximal polyA site. RIP was performed using rat and mouse anti-Elav antibodies from 12-16 hr embryos. Elav binding to the *Elav* proximal polyA site is shown as a positive control. Primers were designed to detect a region in the CDS or a region immediately downstream from the proximal polyA site (EBS). Error bars represent SEM of four separate immunoprecipitation reactions on independently prepared nuclei. n=4. *P* value reflects paired Student’s T test. **F)** RT-PCR experiments show that transfection of S2 cells with Elav induces endogenous expression of *Dscam1-L* (detected using ‘ext’ primers). Anti-Elav western blot confirms Elav overexpression. Actin is shown as a loading control. **G)** Northern blot detecting GFP for S2 cells transfected with a GFP reporter construct harboring the *Dscam1* long 3′UTR (GFP-3′UTR-L). Overexpression of Elav resulted in reduced short and increased long transcripts from the reporter. GAPDH2 Northern is shown as a loading control. L/S represents ratio of long to short 3′ UTR transcripts.

We hypothesized that neurons selectively express *Dscam1-L* given the known neuronal enrichment of Elav and its low or undetectable expression in glia (Berger et al., 2007). To test this, we performed Fluorescence Activated Cell Sorting (FACS) experiments. Using the LexA system (Pfeiffer et al., 2010) we generated flies that simultaneously express mCherry from a glial-specific driver (*repo-Gal4*) and GFP from a neuron-specific driver (*elav-LexA*) (Figure 1C). After FACS sorting of dissected adult brains of these animals, RT-qPCR was performed using primers to detect all *Dscam1* transcripts (“uni”) or *Dscam1-L* transcripts (extension “ext”). *Dscam1-L* was found to be ~20-fold higher in the sorted neurons versus glia, whereas total *Dscam1* mRNA levels were unchanged between neurons and glia (Figure 1C). These data show that *Dscam1-L* expression is more abundant in, if not exclusive to, Elav-positive neurons.

To determine whether Elav regulates biogenesis of *Dscam1-L in vivo*, we performed shRNA knockdown of *elav* and the related gene *found in neurons* (*fne*) in neurons and monitored *Dscam1* 3′ UTR mRNA isoforms by Northern analysis. Elav and FNE have overlapping roles (Zaharieva et al., 2015), and we found that knocking down both genes was lethal in the larval stage (data not shown). Elav/FNE knockdown in late stage embryos (16-20 hr) resulted in a marked decrease in the ratio of *Dscam1-L*/*Dscam1-S*. (Figure 1D).

### Elav binds the *Dscam1* proximal polyA site

Elav has been proposed to compete with the polyadenylation machinery for access to U-rich regions downstream of proximal polyA sites, thus promoting selection of distal polyA sites (Hilgers et al., 2012). Analysis of *Dscam1* 3′ UTR sequence revealed the presence of an U-rich element downstream of the proximal cleavage site. To test whether Elav binds to this motif, we performed Electrophoretic Mobility Shift Assays (EMSAs) and found that Elav bound to a U-rich region downstream of the proximal polyA site in a manner dependent on the integrity of U stretches (Figure S2). To determine whether Elav binds to regions near the proximal polyA site in *vivo*, we performed RNA Immunoprecipitation (RIP) assays on RNA fragments from sonicated 16-20 hr embryo nuclear extracts. Ribonucleoprotein complexes containing Elav were purified using anti-ELAV antibodies followed by RT-qPCR. Elav is known to regulate alternative polyadenylation of its own gene (Hilgers et al., 2012). Thus, as a positive control, we tested for Elav binding to an established elav binding site (EBS) near the proximal polyA site of the *elav* mRNA. By qRT-PCR this region was found to bind Elav (EBS), but not control coding sequence regions (CDS) (Figure 1E). Similarly, Elav binding was observed at the *Dscam1* proximal polyA site (EBS) but not the coding sequence (CDS). Thus, Elav binds at or near the proximal polyA site of *Dscam1 in vivo*.

The Schneider 2 (S2) *Drosophila* cell line does not endogenously express Elav or *Dscam1-L*. Elav overexpression by transient transfection in S2 cells induced expression of *Dscam1-L* (Figure 1F). To determine whether Elav could regulate *Dscam1* long 3′ UTR biogenesis solely via elements in the 3′ UTR, we generated a reporter plasmid containing the *Dscam1* long 3′ UTR sequence, plus U-rich sequence downstream of the distal polyA site, and sub-cloned this downstream of a GFP reporter (*GFP-3′ UTR-L*). Cells transfected with *GFP-3′ UTR-L* predominantly expressed the short 3′ UTR isoform, and only low levels of the long 3′ UTR isoform as measured by Northern blot using a probe against GFP. Overexpression of Elav increased the ratio of long/short reporter mRNA by 3-fold (Figure 1G). Thus, sequences located in *Dscam1* 3′ UTR are sufficient for Elav to promote long 3′ UTR expression.

### Knockdown of *Dscam1-L* in neurons is lethal in adult flies

We next sought out to determine a functional role for *Dscam1-L* transcripts *in vivo*. Using *elav*-GAL4, a neuronal-specific driver, we expressed a shRNA targeting the long (extended) 3′ UTR to neurons (shExt). Northern analysis of dissected male heads showed that *Dscam1-L* was undetectable in the knockdown condition whereas *Dscam1-S* transcripts persisted (Figure 2A). This validated the effectiveness of the shRNA targeting, and also provided additional support that *Dscam1-L* is exclusively expressed in neurons.

**Figure 2:**
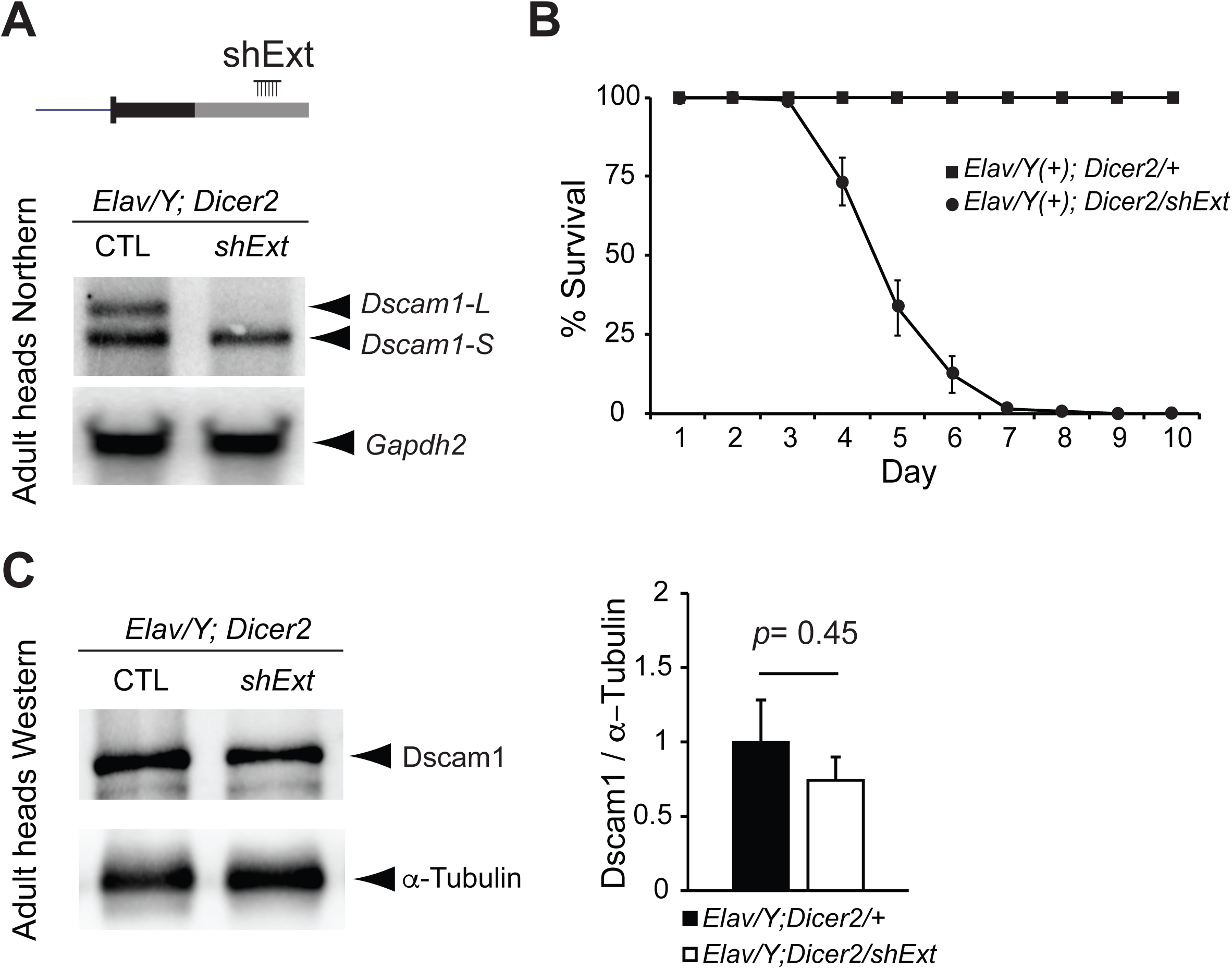
*Dscam1-L* knockdown in neurons is adult lethal. **A)** Top: Schematic showing location of shRNA to knockdown Dscam1-L (*shExt*) using the GAL4-UAS system. Bottom: Northern analysis using a probe for *Dscam1* common exon 11 shows that *shExt* driven by Elav caused loss of *Dscam1-L* in dissected heads. **B)** *Dscam1-L* knockdown caused reduced survival in mixed sex adult flies. **C)** Western analysis of Dscam1 normalized to α-tubulin shows no significant difference in shExt heads compared to control heads. n=7. *P* value represents unpaired Student’s T test. *Elav* represents *ELAV-Gal4*, *Dicer2* represents *UAS-Dicer2*. Also see Video #1.

Neuronal knockdown of *Dscam1-L* using shExt yielded progeny that could survive to adulthood, unlike *Dscam1* null flies (Schmucker et al., 2000). However, these animals displayed severely impaired locomotion and could not fly (**Video #1**). We quantified post-eclosion survival of adults and found that all progeny died by 9 days post-eclosion (Figure 2B). Thus, *Dscam1-L* expression in neurons is required for adult locomotion and survival. We anticipated that the loss of the long 3′ UTR isoform would impact translation of Dscam1. To our surprise, despite the loss of *Dscam1-L* in adult heads, Dscam1 protein levels were not significantly reduced as determined by Western analysis (Figure 2C).

### Long-read sequencing demonstrates that *Dscam1-L* preferentially skips exon 19

We were puzzled by the severity of the locomotion phenotype in *Dscam1-L* knockdown animals despite total Dscam1 protein levels being unaffected. We hypothesized that *Dscam1-L* transcripts might harbor particular protein-coding exons which are not found in *Dscam1-S*. Examination of short-read RNA-Seq tracks at the *Dscam1* locus from different stages of embryonic development revealed an apparent trend of alternative splicing that was coincident with the emergence of the long 3′ UTR in 14-16 hr embryos (Figure 3A). In particular, visualization of splicing events revealed that in later stage embryos there was increased skipping of exons 19 and 23 (Figure 3A). To confirm these trends, we monitored skipping of exons 19 and 23 during embryogenesis by RT-PCR. We found that late stage embryos exhibited increased skipping of exons 19 and 23 compared to early stage embryos (Figure 3B).

**Figure 3:**
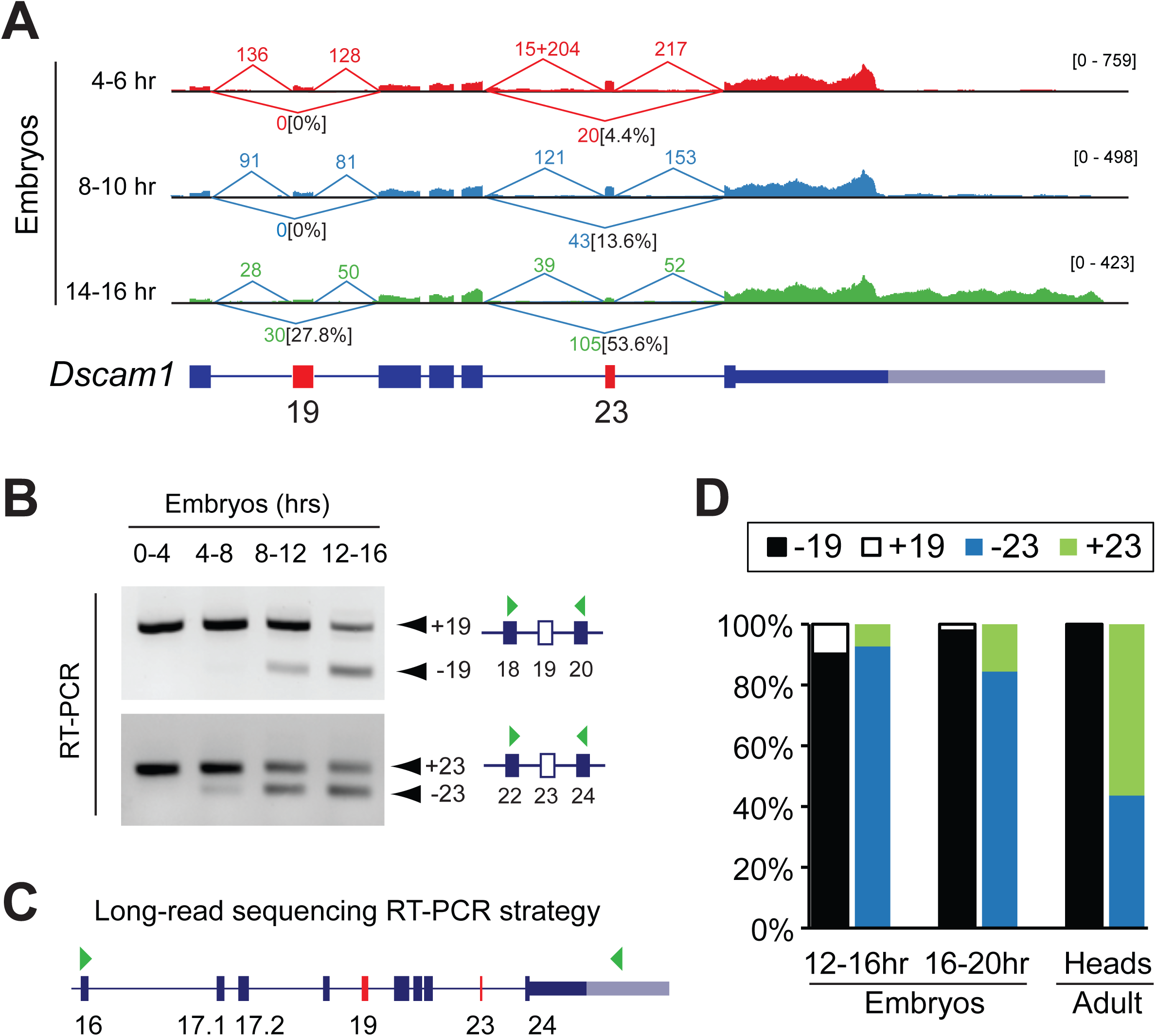
Long-read sequencing demonstrates that *Dscam1-L* transcripts preferentially skip exon 19. **A)** Sashimi plot visualization of RNA-Seq data at the *Dscam1* locus showing spliced read counts from various stages of embryonic development. Note the increased skipping of exon 19 in 14-16 hr embryos that correlates with long 3′ UTR expression. The percentage of reads skipping exons 19 and 23 are noted. **B)** RT-PCR shows skipping of exons 19 and 23 in late versus early stages of embryonic development. **C)** PCR strategy to capture cDNA products from *Dscam1* exon 16 through to the extended 3′ UTR in exon 24 (denoted by green arrowheads) for long read sequencing. Note this strategy detects transcripts exclusively expressing the long 3′ UTR. **D)** Nanopore MinION long read sequencing of RT-PCR products shows a progressive increase of exon 19 skipping during embryonic development, with all adult head *Dscam1-L* isoforms found to skip exon 19. Exon 23 usage in adult heads is mixed for *Dscam1-L*. See also Figure S3.

In order to determine the connectivity of these exon skipping events to the long 3′ UTR, we employed long-read Oxford Nanopore MinION sequencing. Conventional RNA-Seq reads do not provide connectivity information for most transcripts due to the short length of reads. We devised a PCR strategy to capture the full content of *Dscam1-L* from exon 16 through to the long 3′ UTR (Figure 3C). This region contained both constitutively expressed and alternative exons. Note that this strategy provides information only on *Dscam1-L* transcripts, and not *Dscam1-S*. We performed MinION sequencing of these PCR products from cDNA of 12-16 hr embryos, 16-20 hr embryos, and adult heads. Counting of long reads showed that *Dscam1-L* transcripts progressively skipped exon 19 through the time-points. Strikingly, 100% of the reads in adult heads were found to have exon 19 skipped (Figure 3D). In contrast, there was mixed inclusion/exclusion of exon 23 usage for long 3′ UTR transcripts in adult head. This shows the preference for *Dscam1-L* transcripts to skip exon 19 in adult head. Analysis of the nanopore revealed other interesting features of *Dscam1* long 3′ UTR mRNAs (Figure S3). For instance, novel microexons flanking either side of exon 23 were observed in some amplicons, providing additional complexity to *Dscam1* transcript isoforms (Figure S3).

Exon 18 of *Dscam1* harbors additional complexity. Exon 18 uses two distinct splice donor sites that differ by 12 nt (Yu et al., 2009). The slightly longer exon 18 encodes 4 additional amino acids (T-V-I-S) (Figure S4A). Here, we call this variant exon 18-shifted (18s). We analyzed short read RNA-Seq data during embryonic development, L3 CNS, and White pre-pupae (WPP) CNS to determine the isoform usage and connectivity of exons 18, 18s, 19, and 20. For this analysis, we chose short-read RNA-Seq datasets because 150 nt reads were long enough to resolve the connectivity, and they had a lower sequencing error rate compared to the MinION data. This analysis showed that transcripts skipping exon 19 predominantly used exon 18s (18s | −19) versus exon 18 (18 | −19). In contrast, transcripts including exon 19 rarely used 18s (Figure S4B). Thus, in addition to CNS samples preferentially skipping exon 19, they also prefer usage of 18s.

### *Dscam1-L* loss impairs axon outgrowth

To provide conclusive evidence of the connectivity between exon 19 skipped transcripts and the long 3′ UTR, we obtained shRNA knockdown lines that have been previously validated to target *Dscam1* transcripts including exon 19 (sh+19) or excluding exon 19 (sh-19) (Yu et al., 2009) (Figure 4A). The sh-19 shRNA targets the sequence spanning the junction of the exon 18s 5′ splice site, and exon 20 3′ splice site (18s | −19) (Figure S4A) (Yu et al., 2009). sh +19 targeted the sequence spanning exon 18 5′ splice site, and exon 19 3′ splice site (18 | +19).

**Figure 4:**
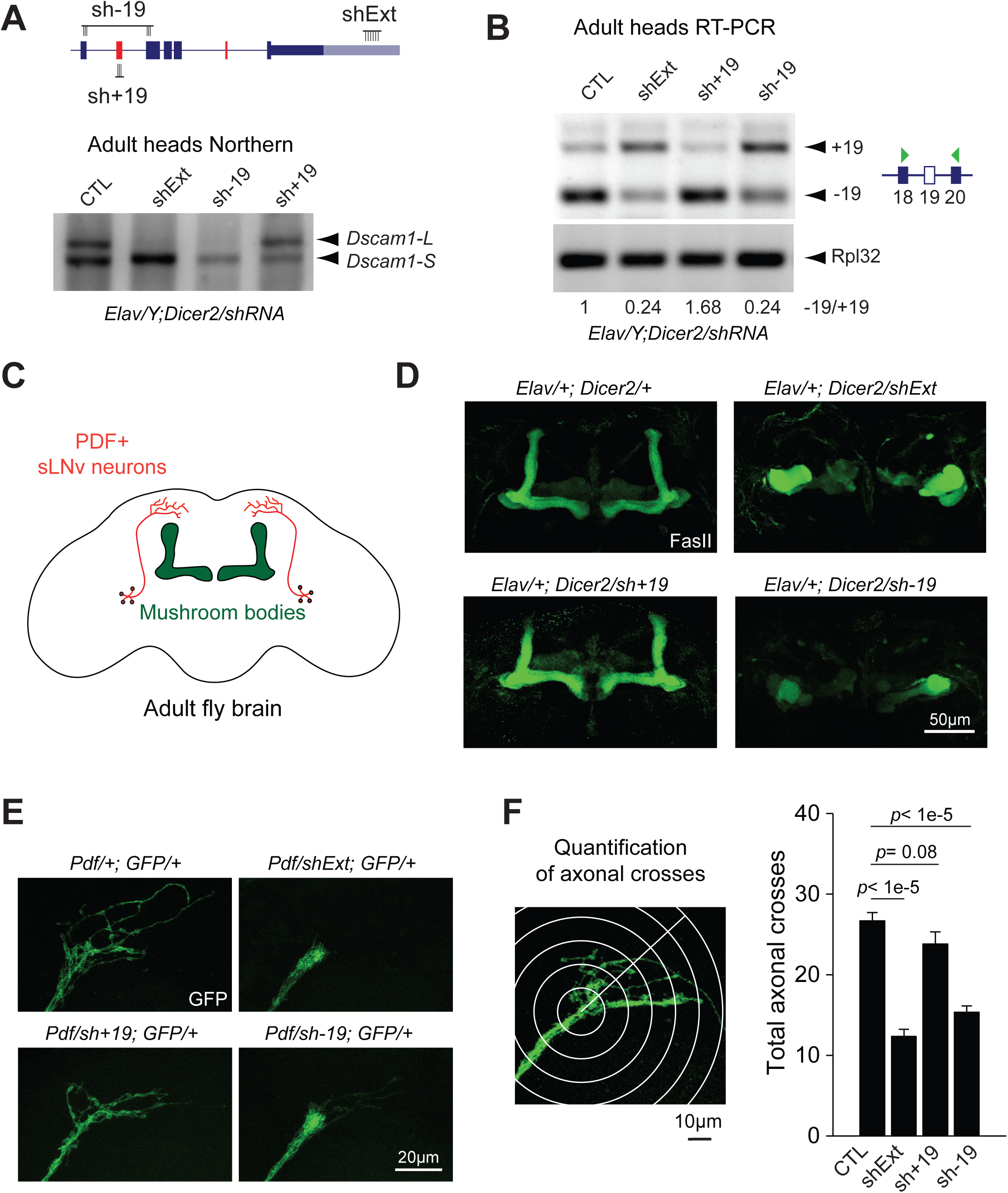
Loss of *Dscam1-L* impairs axon projection. **A)** Top: Schematic of shRNA constructs targeting specific *Dscam1* splice junctions (sh-19, sh+19) and the long 3′ UTR (shExt). Bottom: Northern Blot of male heads from Elav-driven shRNA knockdown shows loss of *Dscam1-L* upon knockdown with sh-19 or shExt, but not sh+19. **B)** RT-PCR shows exon 19 skipping patterns under knockdown conditions in adult heads. **C)** Schematic showing central brain and mushroom body anatomy, with mushroom body highlighted in green and sLNv neurons highlighted in red. **D)** Staining of adult brains with anti-Fasciclin II reveals massive disorganization of the central brain and impaired bifurcation of the mushroom bodies upon neuronal knockdown using shExt and sh-19, but not sh+19. **E)** Impaired axonal outgrowth in adult sLNv neurons expressing sh −19 or shExt driven by *Pdf-Gal4*. **F)** Left: Methodology of axonal outgrowth quantification of PDF positive sLNv neurons. Right: Sholl’s analysis demonstrates that both shExt and sh-19 flies have significantly reduced axonal crossing compared to controls. Error bars represent SEM. n=16-19. *Elav* represents *Elav-GAL4*, *Dicer2* represents *UAS-Dicer2*, *shExt* represents *UAS-shExt*, *sh-19* represents *UAS-sh-19*, *sh+19* represents *UAS-sh+19*, *GFP* represents *UAS-mCD8::EGFP*. See also Videos #1 and #2.

Knockdown in neurons using sh-19 led to effective reduction of *Dscam1-L* as shown by Northern blot of dissected heads confirming that most *Dscam1-L* transcripts skip exon 19 (Figure 4B). As was previously found in the shExt condition, these flies also showed impaired locomotion and could not fly (**Video #2**). In contrast, knockdown using sh+19 did not reduce *Dscam1-L* expression. RT-PCR analysis revealed that knockdown using shExt or sh-19 increased the ratio of +19/-19 in heads (Figure 4C). As expected, sh+19 decreased the ratio of +19/-19 (Figure 4C).

We proceeded to employ these knockdown strategies to examine the cellular functions of *Dscam1-L* transcripts that skip exon 19. Previous work on *Dscam1* null mutants has demonstrated the role of *Dscam1* in axon branching and guidance (Wang et al., 2002). Using anti-Fasciclin II (FasII) staining to image the neuropil in the adult central brain (Figure 4D) we found that neuronal-specific knockdown of *Dscam1-L* using shExt impaired the axonal L-shaped bifurcation structure of the mushroom bodies (MBs), indicating a severe axonal developmental defect (Figure 4E). Neuronal sh-19 knockdown flies showed a very similar loss of MBs bifurcation defect, whereas sh+19 knockdown flies did not.

The above results suggested a role for *Dscam1-L* in axon growth and guidance. To further investigate the nature of this neurodevelopmental impairment, we used *Pdf-GAL4* to drive GFP expression in Pigment Dispersing Factor (PDF) positive ventral lateral neurons (LNv) in adult brain. Among the PDF positive LNvs, the small LNvs (sLNvs) have stereotyped axonal projections toward the dorsal protocerebrum in adult fly brain that can be readily quantified for growth defects (Sivachenko et al., 2013) (Figure 4D). When we knocked down *Dscam1-L* in sLNv neurons, they failed to form arborized axonal terminals (Figure 4F). Using Sholl analysis (Sivachenko et al., 2013), we found that both shExt and sh-19 knockdown in sLNv cells resulted in a significant decrease in the number of axonal crosses, whereas sh+19 knockdown did not (Figure 4F). This defect shows clearly that without exon 19 skipped or long 3′ UTR *Dscam1* mRNAs, neurons cannot undergo proper axonal outgrowth.

### CRISPR deletion of *Dscam1-L*

To validate the importance of *Dscam1-L* for correct axon outgrowth, we used CRISPR/Cas9 to generate flies harboring a deletion of the long 3′ UTR region (*Dscam1*^*ΔL*^). Using a homology directed repair strategy (See Experimental Procedures) we removed the genomic region encompassing the distal polyA site and most of the long 3′ UTR, but retained the proximal polyadenylation site and replaced it with an RFP cassette (Figure 5A). Northern blot of adult heads revealed that *Dscam1-L* was completely lost whereas *Dscam1-S* expression persisted (Figure 5B). As previously found for *Dscam1-L* shRNA knockdown, Dscam1 protein levels in heads were not significantly changed from the wild type condition (Figure 5C). We found that sLNv neurons had reduced axon arborization in *Dscam1*^*ΔL*^ flies (Figure 5D). Impaired bifurcation of the mushroom bodies was also observed, however this phenotype was milder than that observed in the shExt experiments, with <4% of adults showing an abnormally thin or absent lobe (Figure 5E). The milder mushroom body phenotype in the CRISPR *Dscam1*^*ΔL*^ flies (Figure 5E) compared to the shExt condition (Figure 4D) led us to speculate that *Dscam1* −19 transcripts were primarily responsible for mushroom body formation, more so than the long 3′ UTR. To test this, we performed shRNA knockdown of −19 in the *Dscam1*^*ΔL*^ flies (*Elav/+; sh-19, Dscam1*^*ΔL*^/*Dscam1*^*ΔL*^), and found that 100% of these flies failed to properly form mushroom bodies (Figure 5E).

**Figure 5:**
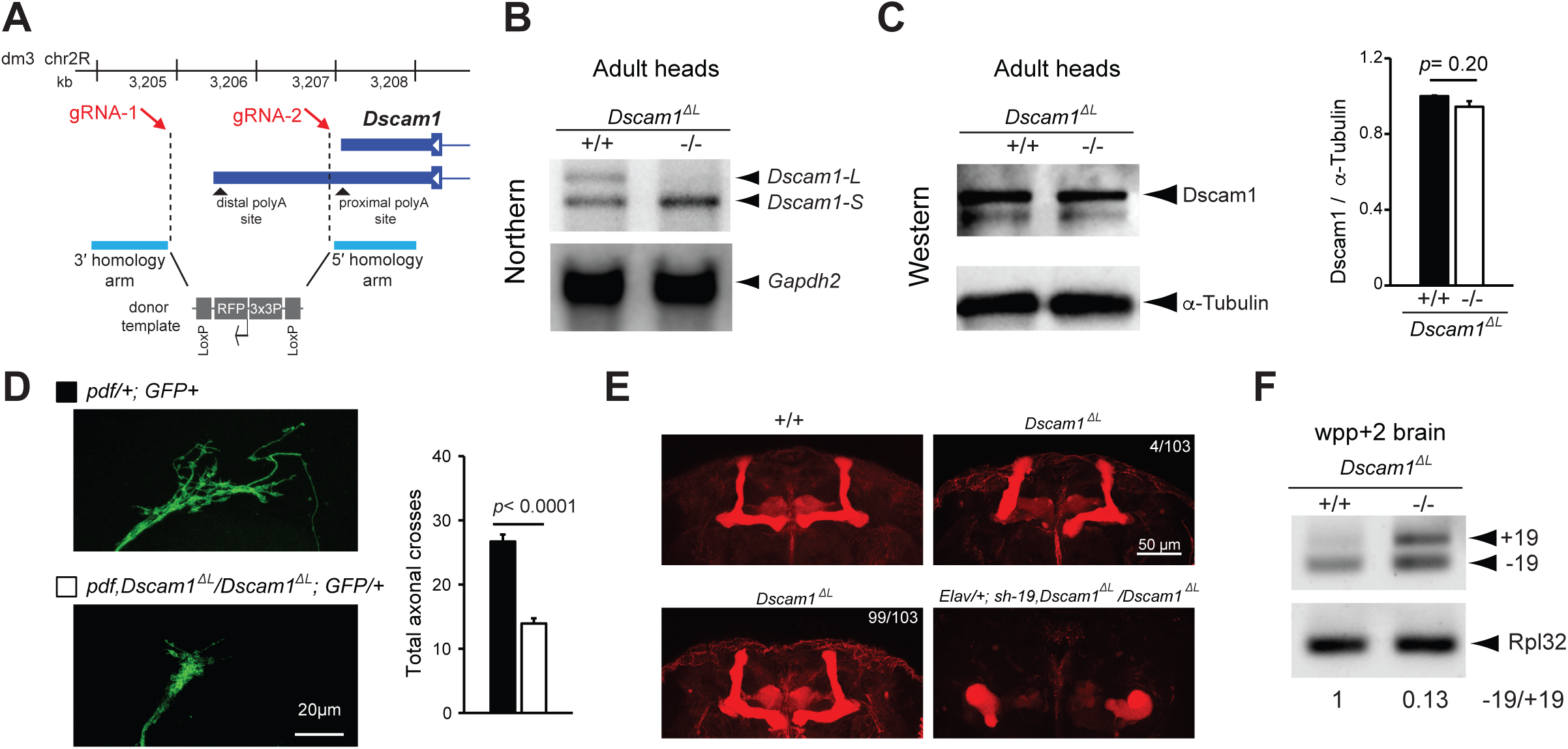
CRISPR-generated *Dscam1*^*ΔL*^ flies show impaired axon projection. **A)** CRISPR/Cas9 genome editing strategy for deletion of the *Dscam1* long 3′ UTR region to generate *Dscam1*^*ΔL*^ flies. Note that the design avoids disruption of the proximal polyA site. **B)** Northern analysis of head lysates shows *Dscam1*^*ΔL*^ flies lack expression of *Dscam1-L*. **C)** Anti-Dscam1 western blot of head lysates and quantification shows that Dscam1 expression is unchanged in *Dscam1*^*ΔL*^ flies compared to control. n=4. **D)** Left side: Representative images of sLNv neuron axon terminals from wild-type and *Dscam1*^*ΔL*^ mutants. Right side: Quantification of sLNv neuron axon terminals shows significantly reduced axonal crosses. *pdf* represents *pdf-GAL4*, and *GFP* represents *UAS-mCD8::EGFP*. See methods for details. n=18-19. All error bars represent SEM. **E)** Staining of adult brains with anti-Fasciclin II reveals impaired bifurcation of the mushroom bodies in *Dscam1*^*ΔL*^ flies. There was low penetrance of this phenotype with 4/103 brains examined showing impaired bifurcation (top right), whereas 99/103 showed normal bifurcation (bottom left). In contrast, knockdown using sh-19 in the *Dscam1*^*ΔL*^ background caused 100% of the flies to have malformed mushroom bodies (bottom right; *Elav/+; sh-19, Dscam1*^*ΔL*^/*Dscam1*^*ΔL*^. 7/7 brains examined). **F)** RT-PCR analysis of white prepupae +2 day brains show increased +19 expression and −19 expression in *Dscam1*^*ΔL*^ mutants.

The increased severity of the mushroom body phenotype in the *Dscam1*^*ΔL*^ flies after knockdown of −19 transcripts suggested that −19 transcripts persist in expression in *Dscam1*^*ΔL*^ flies. Performing a RT-PCR experiment for exon 19 splicing in white pre-pupae brain samples, which are enriched for neurons, revealed that levels of −19 were not reduced, but instead were actually increased (Figure 5F). In addition, there was increased expression of +19 transcripts in the *Dscam1*^*ΔL*^ mutants, which suggested alternative splicing of exon 19 was affected by long 3′ UTR loss (see below). Together, these results suggest that the severe mushroom body phenotype is more attributed to the loss of the −19 transcripts. On the other hand, with no reduction in −19 expression detected in *Dscam1*^*ΔL*^ brains (Figure 5F), the axon outgrowth phenotype in sLNv neurons might be more attributed to loss of the long 3′ UTR.

### *Dscam1* long 3′ UTR is required for regulation of exon 19 skipping

Given the established roles of Elav in alternative splicing, and the connectivity of exon 19 skipping events to the long 3′ UTR, we predicted that Elav might regulate exon 19 skipping. Knockdown of Elav/FNE in 16-20 hr embryos resulted in a decrease in the ratio of −19/+19 as measured by RT-PCR (Figure 6A). S2 cells were found to express the +19 isoform, but nearly undetectable −19 isoform. Transient transfection of Elav led to a moderate increase in −19 expression (Figure 6B). We generated a mini-gene reporter construct to monitor exon 19 splicing that included sequence from exon 18 through to exon 20, including intervening introns. This reporter contained a FLAG tag at the 5′ end to distinguish the reporter cassette from endogenous exon 19 splicing events (Figure 6C). Using RT-PCR, we observed a low level of exon 19 skipping from this reporter construct in the absence of Elav. Strangely, transient co-transfection with Elav failed to promote additional exon skipping (Figure 6D). This suggested to us that the exon 19 skipping event might require additional *cis* elements.

**Figure 6:**
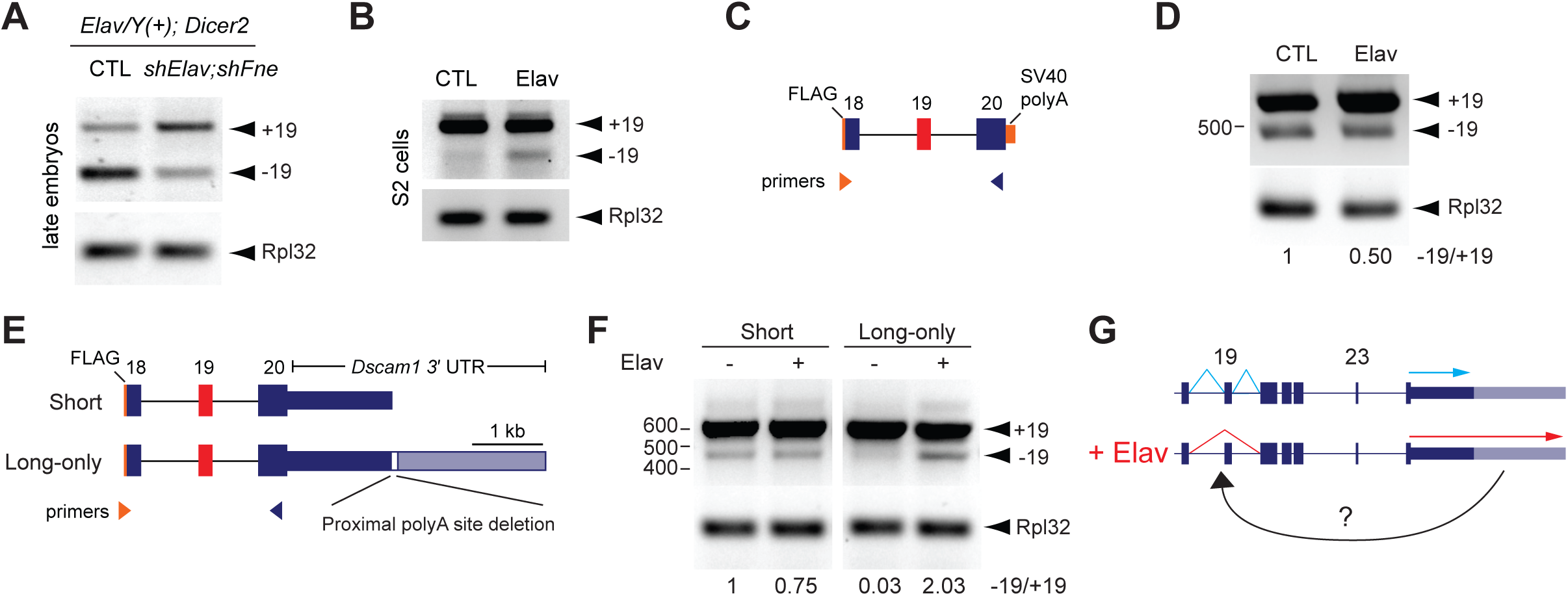
Elav-mediated exon 19 skipping requires the *Dscam1* long 3′ UTR. **A)** RT-PCR analysis of 16-20 hr embryos shows that knockdown of Elav and the related protein FNE by shRNAs in neurons resulted in reduced −19 versus +19 transcripts. *Elav* represents *Elav-GAL4*, and *Dicer2* represents *UAS-Dicer2*. **B)** Elav overexpression in S2 cells induced skipping of endogenous exon 19 as measured by RT-PCR. **C)** Schematic of exon 19 splicing mini-gene reporter. Note Flag sequence at 5′ end allows for distinguishing reporter exon 19 splicing from endogenous exon 19 splicing. **D)** RT-PCR analysis shows no increase in exon 19 skipping from the reporter. **E)** Top: schematic of mini-gene reporters and RT-PCR strategy. Note “long-only” reporter has sequence constituting the proximal polyA site deleted. **F)** RT-PCR analysis shows exon 19 skipping was only induced from the “long-only” reporter upon Elav overexpression.

We next tested whether the presence of the long 3′ UTR influences exon 19 alternative splicing. Mini-gene reporter constructs were generated that harbored the genomic region from exon 18 through exon 20 fused to the *Dscam1* short 3′ UTR sequence (“Short”) or long 3′ UTR sequence (“Long-only”). The “Long-only” reporter had the proximal polyA site deleted in order to prevent short 3′ UTR biogenesis and force expression of the long 3′ UTR. (Figure 6E). No artificial polyA sequences were included at the 3′ end of these reporters. Reporter constructs were co-transfected with Elav in S2 cells and skipping of exon 19 was measured by RT-PCR. The mini-gene reporter harboring the short *Dscam1* 3′ UTR showed mostly inclusion of exon 19, and Elav overexpression failed to induce additional exon skipping (Figure 6F). For the “Long-only” reporter there was less expression of −19 in S2 cells in contrast to the “short” reporter, implying that the long 3′ UTR prevented exon 19 skipping in the absence of Elav. Strikingly, co-transfection of Elav with the “Long-only” reporter led to a clear induction of exon 19 skipping (Figure 6F). Together with the observation that loss of the long 3′ UTR in pupal brain leads to de-regulated alternative splicing of exon 19 (Figure 5F), these results suggest that the long 3′ UTR contains information required for Elav-mediated skipping of exon 19.

## Discussion

APA most commonly changes the length of the 3′ UTR but does not alter protein coding sequence (3′ UTR APA) (Miura et al., 2014; Tian and Manley, 2017). APA can also alter the entire terminal exon resulting in unique 3′ UTRs and C-terminal protein-coding sequences (Alternative Last Exon APA) (Tian and Manley, 2017). Our work here presents a distinct paradigm involving 3′ UTR APA of *Dscam1* in the terminal exon (exon 24) being linked to skipping of a distant upstream exon (exon 19). This coordination of alternative splicing and APA is achieved via co-regulation by the RBP Elav (Figure 6G). This represents a new category of RNA biogenesis regulation in which polyA site choice and upstream alternative splicing events are linked.

Given the connectivity of the *Dscam1* long 3′ UTR to skipping of exon 19, it is difficult to decouple the relative contribution of the long 3′ UTR versus −19 transcripts to the neural phenotypes we observed. Some hints, however, can be gleaned by the differences in phenotype severity between the long 3′ UTR mutants versus the long 3′ UTR knockdown flies. *Dscam1^ΔL^* flies and shExt knockdown flies had similar axon outgrowth phenotypes in sLNv neurons. On the other hand, the mushroom body phenotype was stronger in the knockdown condition compared to the mutant. We observed that −19 transcripts persisted in *Dscam1^ΔL^* pupae nervous system (Figure 5F), whereas −19 transcripts were reduced in the shExt condition (Figure 4B). One interpretation of these differences is that the long 3′ UTR itself is required for sLNv axon outgrowth by conferring mRNA stability, mRNA localization, or translational control, whereas the mushroom body phenotype is primarily attributed to the loss of the protein encoded by −19. In addition, the mushroom body phenotype observed here in the long 3′ UTR shRNA knockdown condition is stronger than that observed when Dscam1 homophilic adhesion is removed (Sawaya et al., 2008), suggesting an axon growth function might also be affected (Kim et al., 2013). Although previous studies have investigated the functional roles of retaining or skipping exons 19 and 23 (Yu et al., 2009), it is unclear how the unique protein sequence of the “-19” isoform impacts Dscam1 binding or activity.

The approaches employed here of shRNA knockdown and CRISPR/Cas9 genome editing can be readily applied to the study of the long 3′ UTR isoforms of other APA regulated genes. Of particular interest, several genes with roles in axon growth and development such as *comm* and *fas1* generate alternative short and long 3′ UTR isoforms (Elkins et al., 1990; Simionato et al., 2007; Smibert et al., 2012). There are hundreds of neural-specific long 3′ UTR isoforms in *Drosophila* alone with unknown functions that remain to be interrogated *in vivo* using these approaches (Hilgers et al., 2012; Smibert et al., 2012).

The specific combination of *Dscam1* long 3′ UTR and alternative exon selection (*Dscam1-L*, −19) is restricted to neurons due to the expression pattern of Elav. Are other *Drosophila* genes regulated in this fashion? Given evidence that Elav can bind to DNA at promoter regions and affect APA (Oktaba et al., 2015), perhaps Elav-regulated APA events are also tied to alternative promoter selection in addition to alternative exon skipping. Do other RBPs with cell-specific expression patterns promote coupling of alternative splicing and APA in other cell types? Incorporating long-read sequencing technology with single cell RNA sequencing workflows could tackle this question on a genome-wide scale. Long-read sequencing has recently revealed a coupling among transcription initiation, alternative splicing and APA in cultured human breast cancer cells (Anvar et al., 2018). As these technologies improve, more insights will surely be gained regarding the genome-wide scope of coordinated alternative splicing and APA.

Our data suggests that a novel function for long 3′ UTRs is regulating alternative splicing of upstream protein-coding exons. Loss of the *Dscam1* long 3′ UTR by deletion conditions was found to de-regulate exon 19 skipping *in vivo* (Figure 5F). Mini-gene reporter experiments showed that the content of exon 19 flanking introns were not sufficient for Elav to promote exon skipping– only when the long 3′ UTR is appended to the reporter can Elav-induced skipping of exon 19 occur (Figure 6F). Further work is needed to identify the precise mechanism that couples *Dscam1* long 3′ UTR selection to skipping of exon 19 (Figure 6G). An intriguing possibility is that Elav binds along the long 3′ UTR while the pre-mRNA is being transcribed, and then is delivered to upstream cis-elements around exon 19 to promote alternative splicing. Such a mechanism might require additional RBPs binding to the *Dscam1* long 3′ UTR, and could involve looping of the pre-mRNA to bring the long 3′ UTR and exon 19 into spatial proximity.

## Acknowledgements

Thanks to Dr. Tzumin Lee for providing *Dscam1* shRNA lines and the Dscam1 antibodies. Thanks to Dr. Kelly Phelps and Dr. Ian Wallace for guidance in preparing purified Elav protein. Thanks to Dr. Valérie Hilgers for sharing RIP RT-qPCR primer sequences. A special thanks to members of the Miura lab for insights and discussion, and to Dr. Hannah Gruner, Dr. Daphne Cooper, and Bong Min Bae for reading and editing of the manuscript. This work was supported by the National Institute of General Medical Sciences (NIGMS) [grant number: P20 GM103650] and the National Science Foundation [grant number: IOS-1656463]. T.K. was also supported by NIGMS [grant number: P20 GM103554]. UNR Fluorescence-Activated Cell Sorting/Flow Cytometry Shared Resource Laboratory was supported by NIGMS [grant number: P30 GM110767].

## Author Contributions

Conceptualization, P.M., Y.Z., J.K., T.K.; Methodology, P.M., T.K., Z.Z., R.P., K.S., M.B.; Investigation, P.M., T.K., Z.Z., R.P., K.S., M.B., H.N.; Data Curation, P.M., Z.Z., M.B., K.S.; Software, M.B.; Writing – Original Draft, P.M. Z.Z.; Writing – Review & Editing, P.M., Z.Z., T.K., J.K., Y.Z.; Supervision, P.M., T.K.; Funding Acquisition, P.M., J.K., T.K.

## Declaration of Interests

The authors declare no competing interests

## STAR Methods

### Fly breeding and stocks

All flies are raised at 25°C, 12-hr dark/12-hr light cycles on standard food (231g cornmeal, 96 g yeast, 54 g agar, 231 mL molasses, 36 mL Tegosept, 24 mL propionic acid for 6L of food). Elav-GAL4; UAS-Dicer2 was a gift from Eric Lai (Sloan Kettering Institute). Stocks from Bloomington Drosophila Stock Center (BDSC) are: repo-GAL4, (BL#7415), Elav-lexA (BL#52676), 20XUAS-6XmCherry (BL#52267), 13XlexAOP-GFP (BL#32203), and UAS-shElav (BL#28371). UAS-shFne was from Vienna Drosophila Resource Center (VDRC, # 101508).

### RNA-Seq track visualization

Visualization of RNA-Seq. bam files generated by the modENCODE consortium was performed using Integrated Genomics Viewer (Thorvaldsdottir et al., 2013). RNA-Seq tracks used for visualization of 3′ UTR extension of *Dscam1* included: L3 CNS (SRR070410), 6-8 hr after egg laying (a.e.g.) embryos (SRX246422), 16-18 hr embryos (SRX246418). RNA-Seq tracks used for sashimi-plot splicing visualization included 4-6 hr embryos (SRX246408), 8-10 hr embryos (SRX246409), and 14-16 hr embryos (SRX246412). Sashimi plots were generated in IGV using minimum exon coverage “7”.

### Western Analysis

Fly heads or embryos were collected and lysed with protein extraction buffer (1% Triton X-100, 100mM pH 6.8 Tris-HCl, 150mM NaCl, and protease inhibitor tablet (Thermofisher Scientific A32955)). Protein samples were prepared with denaturing buffer (5% β-Mercaptoethanol, 0.2M pH6.8 Tris-HCl, 8% SDS, 40% Glycerol, 0.1% Bromophenol blue) and boiled at 95°C for 5 mins. Fifteen μg of sample was loaded in each well of a 7% SDS-PAGE gel, and then transferred to the membrane with Turbo Mini PVDF Transfer Pack (#1704156, Bio-Rad). The blot was then blocked with 5% BSA in TBST (0.5% Tween-20 in TBS), incubated with primary antibody at 4°C overnight, washed with TBST and incubated with secondary antibody (Jackson ImmunoResearch Inc.) at room temperature (RT) for 1 hour, and washed prior to ECL detection (20-302B, Genesee Scientific Corporation) and imaged using the ChemiDocTM Touch Imaging System (Bio-Rad Laboratories, Inc). Rabbit anti-DSCAM-cytoplasmic domain was used at 1:1500 (gift from Dr. Dietmar Schmucker) (Dascenco et al., 2015). Anti-alpha-tubulin at was used at 1:400 (12G10, Developmental Studies Hybridoma Bank), rat anti-Elav at 1:750 (7E8A10, Developmental Studies Hybridoma Bank), and anti-actin at 1:100 (JLA20, Developmental Studies Hybridoma Bank).

### Immunohistochemistry

Flies were collected and brains dissected in phosphate-buffered saline (PBS), and then fixed in 4% PFA at RT for 30 mins, blocked with 5% normal goat serum (NGS) in PBST (0.3% Trition X-100 in PBS) at RT for 1 hr. Samples were incubated with primary at 4°C overnight, and washed 3×15 min in PBST. Secondary antibody incubation (Jackson ImmunoResearch Inc.) was performed at R.T. for 1 hour, then washed 3 × 15 min with PBST and mounted with Vectashield Mounting Medium (H-1000, Vector Laboratories). Imaging was performed using a Leica TCS SP8 microscope. Anti-Fasciclin II was used at 1:20 (1D4, Developmental Studies Hybridoma Bank). Anti-mouse Cy3 secondary antibody was used at 1:500 (Cat# 515-165-003, Jackson ImmunoResearch).

### S2 Cell Transfection

Schneider 2 cells were obtained from Drosophila Genomics Resource Center (DGRC). Cells were cultured in Schneider’s *Drosophila* medium (ThermoFisher CAT# 21720024), with the addition of 10% fetal bovine serum (Atlanta Biologicals CAT# S11150). For transient transfection, cells at 80% confluent were transfected using Effectene transfection reagent (Qiagen CAT# 301425) according to the manufacturer’s protocol for suspension cells. For co-transfections, 600 ng of Ubiquitin-GAL4 plasmid (gift from Eric Lai), 300 ng of reporter, and 300 ng of UAS-ELAV or empty vector control was added (per 35 mm well). Cells were collected 48 hours post-transfection and pelleted in preparation for RNA extraction. UAS-Elav contained the coding region of Elav with sequence encoding for FLAG-HA tag at the N-terminus (5′-GACTACAAGGACGACGATGACAAGTACCCTTATGACGTGCCCGATTACGCT-3′) and was cloned using pUASTattB vector (gift from Eric Lai).

### RNA Extraction

RNA extracted from cell culture and *Drosophila* was performed using TRIzol reagent (Invitrogen CAT# 15596026) following the manufacturer’s protocol. Following extraction, and prior to cDNA synthesis, RNA was DNase treated using the DNA-*free* DNase Treatment and Removal Reagents (Ambion CAT# 1906) following the manufacturer’s “Routine DNase treatment” protocol.

### cDNA Preparation, qPCR, end-point PCR

DNase treated RNA was reverse transcribed into cDNA using iScript Reverse Transcription Supermix for RT-qPCR (Bio-Rad CAT# 1708840) as ^per manufacturer’s instructions. cDNA was then diluted 1:5 in ddH_2_O. Real-time PCR was^ performed using SYBR Green PCR Master Mix (Invitrogen CAT# 4309155). Real time PCR was performed using a CFX96 Touch Real-Time PCR Detection System (Bio-Rad CAT#1855195). All data was analyzed using Bio-Rad CFX software. End point PCR was performed using Taq polymerase at optimized cycle number, and products were run on 1-2.5% Agarose gels with EtBr prior to imaging. Primer sequences can be found in table S1.

### Northern Analysis

Northern analysis was performed as previously described (Gruner et al., 2016) using P^32^ labeled DNA probes.

### *Dscam1* 3′ UTR Reporter Cloning

The long *Dscam1* 3′ UTR starting from after the stop codon until past the distal poly A site was amplified using iProof High-fidelity PCR kit (Bio-Rad Cat#1725330) and subcloned into GFP reporter constructs plasmids (Kim et al., 2013) from which the SV40 polyA site was removed.

### *Dscam1* Mini-gene Reporter Cloning

The sequence of *Dscam1* from exon 18 to 20 was amplified using LongAmp Taq polymerase (FLAG tag was added to the 5′ end) and subcloned into the above GFP reporter. The GFP coding sequence was removed by restriction digest.

Dscam1 short or Dscam1 long-only was subcloned downstream of exon 20. For the “Long-only” 3′ UTR, we introduced a 64-nt deletion (AAATATATGAT TTTGATTTTATTTTTAATTGATTACGTTCGCTTTTGTTTGATTATTGTTTTGG) to remove the proximal polyA site using HiFi Assembly Cloning Kit (NEB CAT#E2621).

### RIP RT-qPCR

Embryos (12-16hr) were collected and nuclear extraction was performed following a previously described protocol (Hilgers et al., 2012). Nuclei were sonicated for 10 cycles of 30-sec on/30-sec off on Branson Sonifier 450. Nuclear extracts containing fragmented RNA were subjected to immunoprecipitation following the standard protocol of Dynabeads Protein G Immunoprecipitation Kit (CAT#1007D). Samples were incubated overnight at 4°C with a mixture of 1 μg rat and 1 μg mouse anti-Elav antibodies (DSHB CAT#7E8A10 and CAT#9F8A9, respectively) or a mixture of 1 μg rat and 1 μg mouse IgG (Rockland CAT#012-0102 and CAT#010-0102, respectively). Proteinase K treatment was performed prior to reversal of of cross-links performed for 1 hr × 68 °C. RNA was extracted from input control immunoprecipitates using Trizol and then treated using Turbo DNA free Kit. After performing reverse transcription using random hexamers, samples were used at 1:5 dilution for qPCR.

### Generation of shRNA lines

For generating the shExt shRNA construct the following primers were used: Forward: 5′-GATCGCTAGCAGTAGGCGTTTAGTTTCACTTCAATAGTTATATTCAAGCATA-3′; Reverse: 5′-GATCGAATTCGCAGGCGTTTAGTTTCACTTCAATATGCTTGAATATAACTA-3′. NheI and EcoRI were the restriction sites used to clone the short hairpin into pWALIUM20 vector (Harvard Medical School). Plasmid DNA was obtained using QIAGEN Plasmid Plus Midi Kit (Qiagen CAT#12945) and sent for microinjection (BestGene) into the y1 w67c23;P{CaryP}attP40 strain. Other shRNA lines including sh-19, sh+19 have been previously described (Yu et al., 2009) and were gifts from Dr. Bing Ye (University of Michigan).

### Generation of *Dscam1*^*ΔL*^ flies by CRISPR/Cas9 Deletion

CRISPR/Cas9 genome editing was performed by Well Genetics. Two gRNAs were designed flanking the extended 3′ UTR of *Dscam1* which caused double-stranded breaks at positions chr2R: 3,204,852 and chr2R: 3,206,926 generating a nearly 2 kb deletion (dm3 genome coordinates). A 134 bp region between the proximal *Dscam1* short 3′ UTR cleavage site and the first double-stranded break was left intact in order to not disturb any potential DSEs important to proper cleavage and polyadenylation of the short 3′ UTR isoform. Homology directed repair was used to knock-in a 1.8 kb RFP cassette containing loxP sites. RFP was used as a visible marker for screening successful mutants. Three successful *Dscam1^ΔL^* mutants were generated with the expected deletion and knock-in. Flies were balanced over CyO or GFP, CyO to create stable stocks.

### Longevity Assay

Flies were collected within 24 hr after eclosion, and raised 20 flies per vial (male and female) with standard food. The number of surviving flies was recorded every day at same time point.

### Embryo Collection

Flies with the genotype of interest were placed in cages containing grape agar and thin layer of yeast paste to lay eggs. Flies were synchronized by changing the plate two times after over the course of 24 hours. Following synchronization, flies were allowed to lay eggs for 4 hours per plate. Plates were then allowed to develop to the required time points. Embryos were dechorionated with 50% bleach and rinsed with water prior to downstream RNA or protein analysis.

### Sholl Analysis

1 to 3 days old male flies were collected, and kept for two days before dissection. Male brains were dissected and fixed in 4% PFA (paraformaldehyde) in PBST for 30 mins at room temperature, washed 3×15 min with PBST, and then mounted with Vectashield Mounting Medium. Samples were imaged under Leica TCS SP8 microscope, and then images were quantified with Image J sholl analysis plugin (Fernandez et al., 2008). Six concentric rings (radius step size=10 μm) centered at the point where the first-class branches open up were drawn on each brain hemisphere. The total number of intersections of all branches crossing the six rings was counted. Data was collected from both brain hemispheres and the average of them was used as one sample for quantification. For statistical test, Mann Whitney test for non parametric samples was used.

### FAC sorting of neurons and glia from adult brains

To generate a fly expressing fluorescence tags for neurons and glia, *Elav-lexA* (BL#52676), *20XUAS-6XmCherry* (BL#52267), *13XlexAOP-GFP* (BL#32203), and *repo-GAL4* (BL#7415), and *y*^1^*w*^1^;*CyO/Sco;MKRS/Tm6B,Tb* were used to generate *Elav-lexA/20XUAS-6XmCherry;13XlexAOP-GFP/repo-GAL4*. All of these stocks were balanced using *y*^1^*w*^1^;*CyO/Sco;MKRS/Tm6B,Tb*. Preparation of brains for FACS was performed based on a previously described protocol (Nagoshi et al., 2010). Briefly, 40 – 50 adult brains were dissected in a dissection saline (9.9 mM HEPES-KOH buffer, 137 mM NaCl, 5.4 mM KCl, 0.17 mM NaH_2_PO_4_, 0.22 mM KH_2_PO_4_, 3.3 mM glucose, 43.8 mM sucrose, pH 7.4) containing 50 μM D(–)-2-amino-5-phosphonovaleric acid (AP5), 20 μM 6,7-dinitroquinoxaline-2,3-dione (DNQX), 0.1 μM tetrodotoxin (TTX) and then transferred to a modified SMactive medium (SMactive medium containing 5 mM Bis-Tris, 50 μM AP5, 20 μM DNQX, 0.1 μM TTX). After dissection, brains were washed again with the dissection saline and then digested with L-cysteine-activated papain (50 units/mL in dissection saline; Worthington). Reaction was then quenched with SMactive media and the brains were triturated with a flame-rounded 1000-μL pipette tip and a flame-rounded 200-μL pipette tip. Prior to FACS selections, Hoechst 33528 was added as a viability marker. Individual samples were sorted for neurons (*Elav*-positive cells) and glia (*repo*-positive cells) using the BD FACSAria II SORP with 70 μm nozzle at 70 psi. FACS was performed at the Fluorescence-Activated Cell Sorting/Flow Cytometry Shared Resource Laboratory at the UNR School of Medicine. RNA from isolated neurons and glia was extracted using Trizol.

### Nanopore Minion sequencing

Amplification from cDNA of a region encompassing Dscam1 exon 16 until within the extended 3′ UTR was performed using the following primers (forward: 5′TTTCTGTTGGTGCTGATATTGCCGAATACGACTTTGCCACCT-3′; reverse: 5′-ACTTGCCTGTCGCTCTATCTTCTCTGTAGCTCCATTGCATCG-3′). The PCR product was purified using AMPure XP beads (A63881, Beckman Coulter Life Science), and then used as a template for PCR amplification using nanopore barcodes primers (SQK-LSK108 Ligation Sequencing Kit 1D, and BC001, BC002, BC010 in EXP-PBC001 PCR Barcoding Kit I, from Oxford Nanopore Technologies) to obtain barcoded amplicons. PCR products (1μg in total) extracted from magnetic beads was used for Nanopore sequencing with FLO-MIN 107 flowcell. Raw sequence reads are deposited at the sequence read archive (https://www.ncbi.nlm.nih.gov/sra). 12-16 hr embryos: SAMN11457160; 16-20 hr embryos: SAMN11457161; adult heads: SAMN11457162.

### Analysis of Minion data

Raw data demultiplexing and basecalling were performed with Albarcore 2.0.1 provided by Oxford Nanopore. Reads were trimmed using porechop version 0.2.3 (https://github.com/rrwick/Porechop) using default settings. Reads were aligned to the full Dscam1 gene sequence from the DM6 assembly using gmap version 2018-02-12 and the ‘-f samse --sam-extended-cigar’ settings. Alignment isoforms were identified and counted using isocount 1.1.0 (https://github.com/bauersmatthew/isocount) and the ‘--antisense -c 75,15’ settings. Based on the overall coverage distributions for each feature, a >75% coverage cutoff was used for feature inclusion and a <15% coverage cutoff was used for feature exclusion. *Dscam1* exon, intron, and 3′ UTR positions were obtained from the Ensembl Dme v89 annotation. Isoforms not conforming to all of the following requirements were excluded from further analysis: 1-Inclusion of both the extended and universal 3’ UTR. 2-Exclusion of all intron sequence. 3-Inclusion of the constitutive exons 16, 18, 20, 21, and 22. 4-Usage of exactly one of exon 17.1 or 17.2.

### Elav Protein Expression and Purification

The coding region of ELAV was PCR amplified from cDNA isolated from w1118 *Drosophila* using AccuPrime Pfx DNA polymerase and subcloned into the pENTR/D-TOPO vector (Invitrogen CAT# K240020) donor vector. The destination vector chosen was pET-60-DEST (Novagen CAT# 71851-3). The destination vector was grown up in One Shot *ccd*B Survival 2T1^R^ cells (Invitrogen CAT# A10460). Both vectors were then prepared using midi-prep (Qiagen CAT# 12945). The final expression vector was made using LR Clonase II (Invitrogen CAT#11791-020) enzyme per the manufacturer’s protocol, which flipped the Elav coding region into the pDEST60 vector with the N-and C-terminal tags. BL21(DE3) Competent E. coli (New England Biolabs CAT# C2527l) were transformed with pDEST60-Elav construct according to the manufacturer’s protocol. Cells were collected by centrifugation in 50 ml centrifuge tubes for 15 minutes at 3000 rpm and resuspended in 5ml of water and centrifuged again to rinse off leftover media. The pellet was resuspended in 30 ml of a buffer containing 25 mM Tris HCl pH 7.5, 150 mM NaCl, 20 mM imidazole and 10 mM MgCl_2_ with the addition of a protease inhibitor tablet (Thermo Scientific CAT#88666) and 30 μg of lysozyme (Sigma CAT# L6876). The mixture was left to incubate for 30 minutes at room temperature. The cells were then sonicated (Branson Sonicator 450) on ice six times for 15 second intervals at a power level of four. The sample was then centrifuged at 28,000 rpm for 1 hour at 4°C. Following sonification, 900 ml of a buffer containing 25 mM Tris HCl pH 7.5, 150 mM NaCl, 20 mM imidazole and 10 mM MgCl_2_ and 290 μl of 2-mercaptoethanol was prepared. 2 ml of glutathione agarose (Thermofisher CAT# 16100) was then mixed with 48 ml of the buffer just prepared, and was allowed to sit so the beads settled. Supernatant was then removed leaving only the beads at the bottom of the tube. The supernatant from the 1-hour centrifugation step was then added to the beads and incubated for 2 hours at 4°C with rocking. The rest of the purification took place at 4°C in a cold room. A chromatography column (Bio-Rad CAT# 7374021) was rinsed with the imidazole buffer containing the 2-mercaptoethanol, and all 50 ml of the supernatant-glutathione bead mix was added to the column, allowing the beads to compact in the column. Some flow-through was kept for quality analysis. To remove any material not bound to the agarose beads, 150 ml of the buffer containing 2-mercaptoethanol was added to the column to wash the beads, and the flow-through was discarded. To remove the Elav bound protein, 0.28g of reduced L-glutathione (Sigma CAT# G4251) was added to 30ml of the buffer containing 2-mercaptoethanol. Ten ml of the reduced L-glutathione solution was added to the column and left to incubate for 5 minutes. Following incubation, 8×1 ml aliquots were collected. To determine which fraction contained the highest concentration of Elav, 5 μl from each fraction (and also the flow-through samples collected) was aliquoted and added to new tubes each containing 5 μl of SDS loading buffer. The samples were heated at 95°C for five minutes and then all samples were ran in a tris glycine gel for 30 minutes. The gel was then stained for 30 minutes while shaking in a solution containing 3.75 ml of acetic acid, 46.25 ml of ddH_2_O and 10 μl of Sypro orange (ThermoFisher CAT# S6650). The gel was then scanned on a gel imager (GE Healthcare Typhoon Trio) to determine which fraction contained the most protein. The fraction with the most protein was quantified using a BSA standard curve protocol, using ImageJ to quantify pixel intensity of the Flamingo-stained (Bio-Rad CAT#161-0490) TrisGlycine Gel.

### EMSA RNA Probe preparation

RNA oligo nucleotide probes was synthesized by Integrated DNA Technologies. See Figure S2 for EBS and Mutant probe sequences. Oligos were then purified by PAGE prior to use. Each 2.5 pmol sample was 5′ labeled with γ^32^-P. Sample was dried by SpeedVac. A master mix containing 1ul of T4 Polynucleotide Kinase buffer, 5.6 μl Water, and 0.4 μl of T4 polynucleotide kinase was prepared. The dried RNA was resuspended with 7 μl of the master mix. Three μl of [γ^32^-P] ATP was added to each sample and left to incubate at 37**°**C for one hour. Probes were then purified using Illustra Microspin G-25, and PAGE purified.

### EMSA

A 4% acrylamide:bisacrylamide 80:1 native gel was prepared (27.25 ml DEPC treated water, 1.65 ml 10×TBE, 3.3 mL 40% acrylamide, 830 μl 2% bisacylamide, 133 μl of 10% APS, and 66.6 μl of Temed) and left to polymerize for at least 2 hours. RNA samples were resuspended in hybridization buffer (10 mM Tris HCl pH 7.5, 100 mM NaCl, and 0.1 mM EDTA) so that the final concentration was 10 times less than the molar concentration of the ELAV protein sample. Samples were then incubated at 65**°**C for 5 minutes. A master mix was prepared consisting of 1 μl 10× reaction buffer (450 mM Tris HCl pH 7.5, 5 mM NaCl, and 400 mM KCl), 1 μl of 250 μg/ml tRNA, 1 μl 5mM DTT, 1 μl 500 μg/ml BSA, and 1 μl of 6 units/μl RNAse inhibitor. Five μl of the hybridization mix was aliquoted per sample and 4.35 μl of ELAV protein, or GST buffer without protein was added to each tube. Then 0.65 μl (154 nM) of RNA sample was added to its respective tube. Hybridization was allowed to take place for 20 minutes at room temperature, and then 3 μl of loading buffer was added to each sample. The gel was pre-run for 15 minutes at 250V, then samples were loaded and run for 140 minutes in a cold room at 4**°**C. Gels were then dried on a gel dryer at 80**°**C for 1 hour with vacuum. A phosphor screen was exposed to the gel overnight and imaged the following day.

---

**Figure S1.**
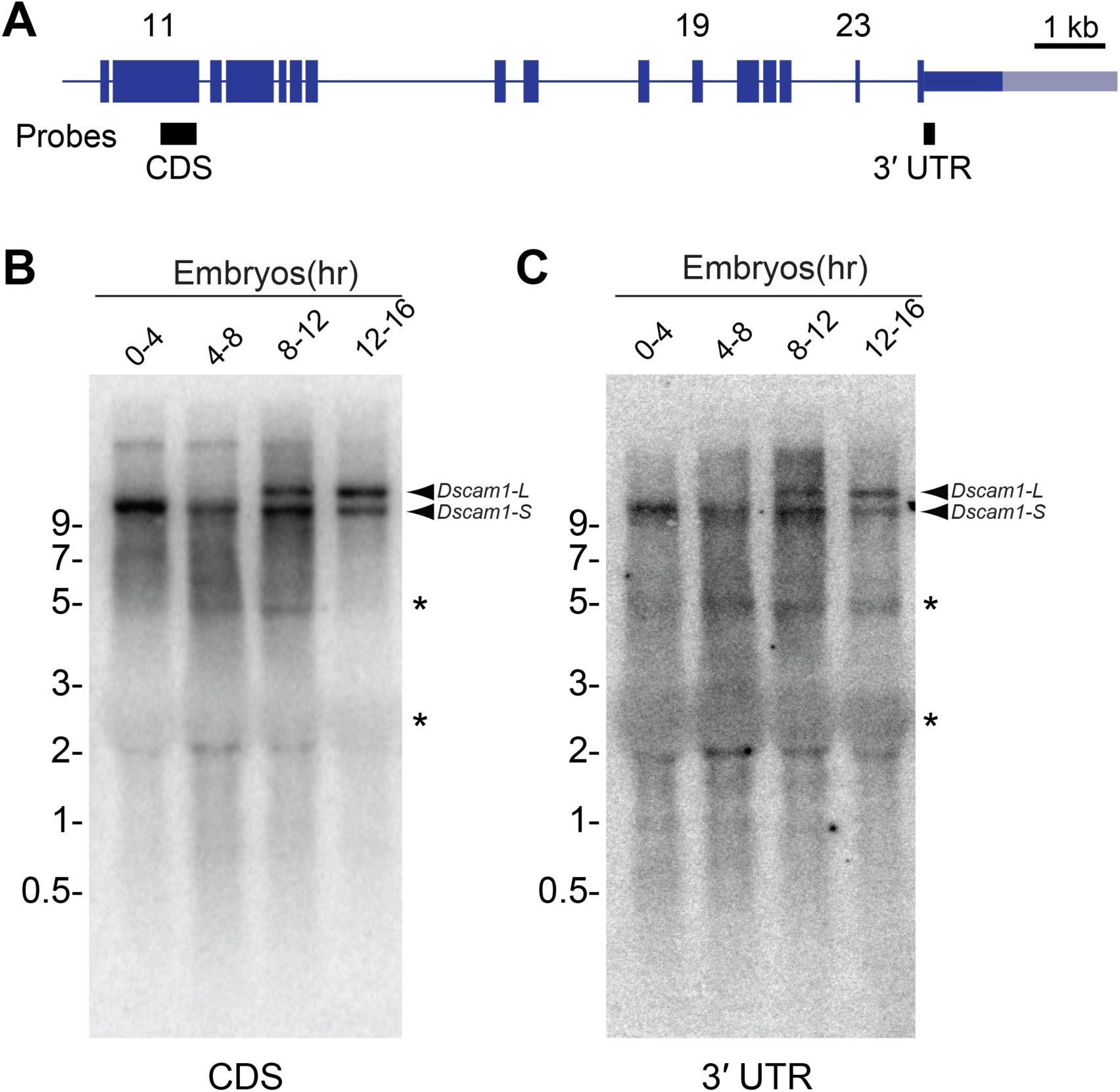
Northern analysis of *Dscam1* during embryonic development. Related to Figure 1. **(A)** Schematic of Northern probe locations. **(B)** Non-cropped northern blot using a probe to *Dscam1* exon 11 which is expressed in all *Dscam1* isoforms (CDS). **(C)** Stripped and re-probed blot using probe targeting the proximal 3′ UTR (3′ UTR). *, denotes common background bands often observed with these Drosophila northern blots.

**Figure S2.**
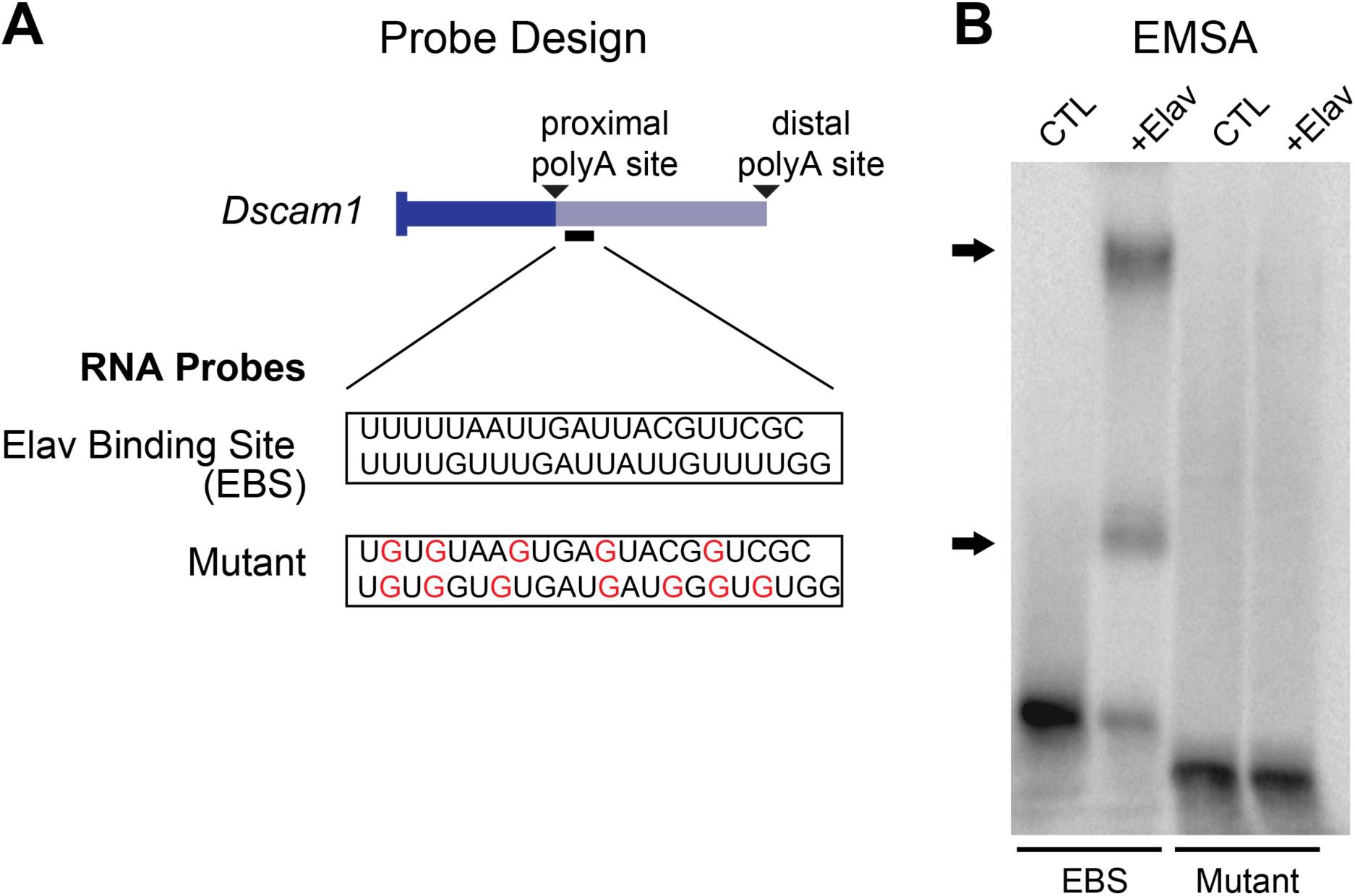
EMSA demonstrates Elav binds a U-rich region downstream of the *Dscam1* proximal polyA site. Related to Figure 1. **(A)** Schematic showing probe design with location of proximal 3′ UTR Elav binding site (EBS) probe used in RNA electrophoretic mobility shift assays (EMSA). Mutant probe has disrupted integrity of U-rich stretches generated by U to G mutations (indicated in red text). **(B)** EMSA reveals that upon the addition of recombinant purified Elav, an upward shift occurs, indicating Elav binding. Elav has been shown to bind as a multimer (Soller and White, 2005) which might explain the higher molecular weight supershift. Mutant probe was incapable of Elav binding, as evidenced by the lack of shift.

**Figure S3.**
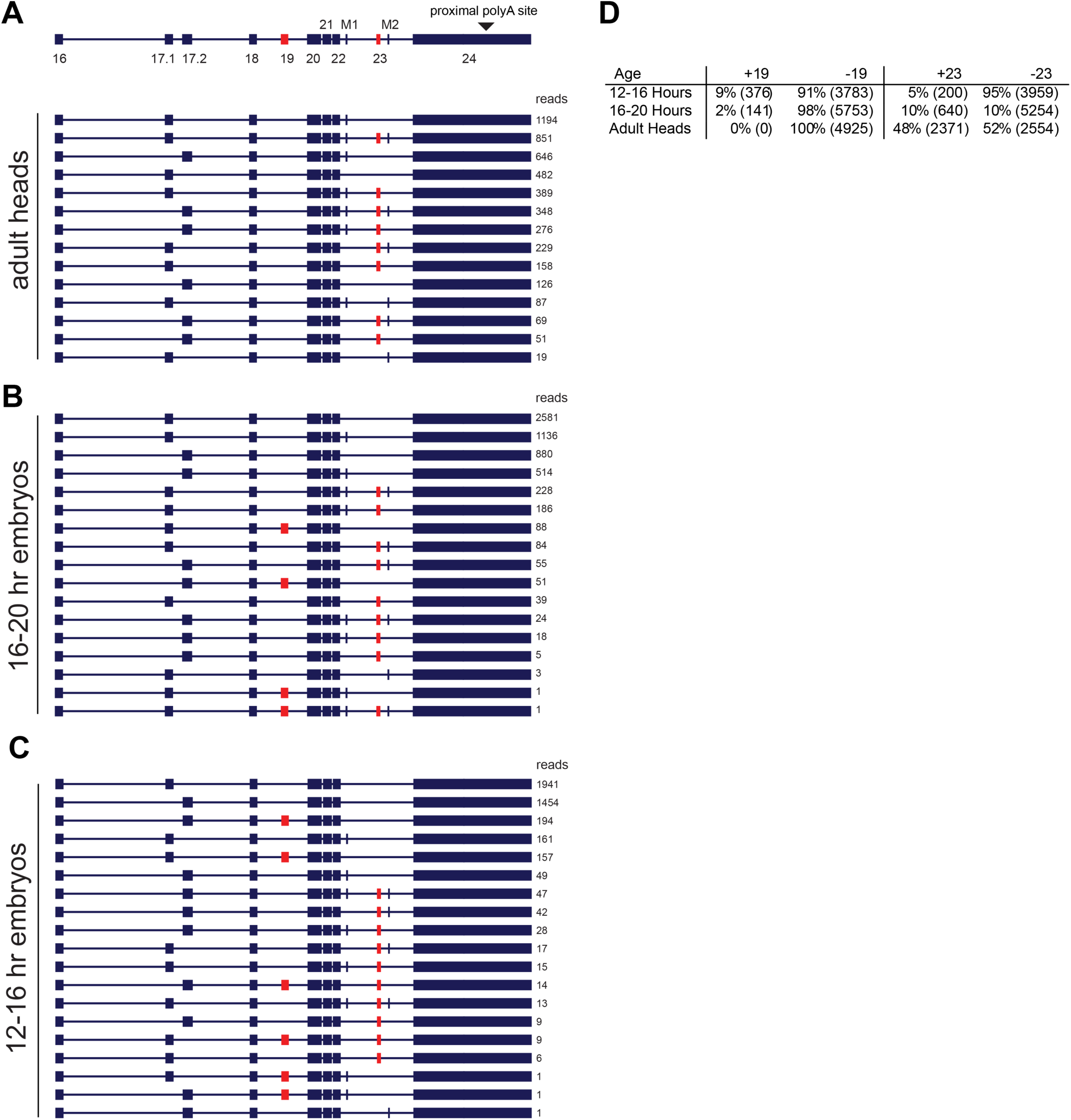
Frequency of *Dscam1* transcript isoforms connected to extended 3′ UTR determined by Nanopore sequencing of *Dscam1*. Related to Figure 3. **(A)** Adult heads. **(B)** 16-20 hr embryos. **(C)** 12-16 hr embryos. **(D)** Summary table showing the frequencies of exons 19 and 23 in transcripts including the extended 3′ UTR. M1-microexon 1, M2-microexon 2.

**Figure S4.**
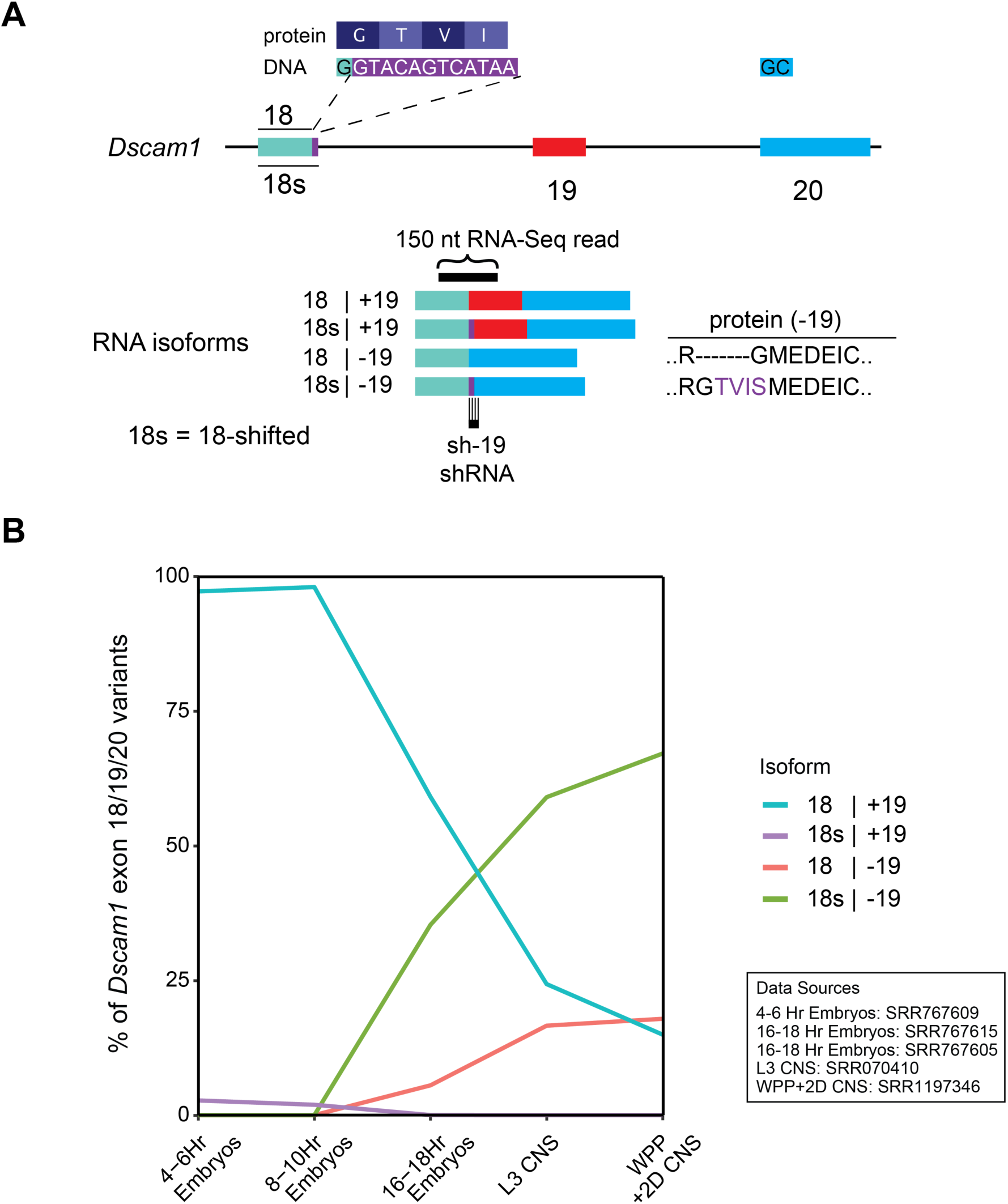
Exon 18/18s/19/20 usage determined from short read RNA-Seq data. Related to Figures 3 & 4. **(A)** Schematic of *Dscam1* exon usage from exons 18 to 20. An exon 18 variant that is 12 nt longer at the 3′ end is named 18-shifted (18s) (highlighted in purple). Compared to the 18 | −19 isoform, the 18s | −19 isoform gains the amino acid sequence T-V-I-S. Note the location of the sh-19 shRNA that uniquely knocks down 18s | −19 isoforms. RNA-Seq short-reads (150 nt) had sufficient length to resolve the connectivity between exons18, 18s, 19, and 20. **(B)** Exon usage during embryonic development, L3 CNS, and WPP +2D CNS as analyzed from short read RNA-Seq data.

**Table S1:**
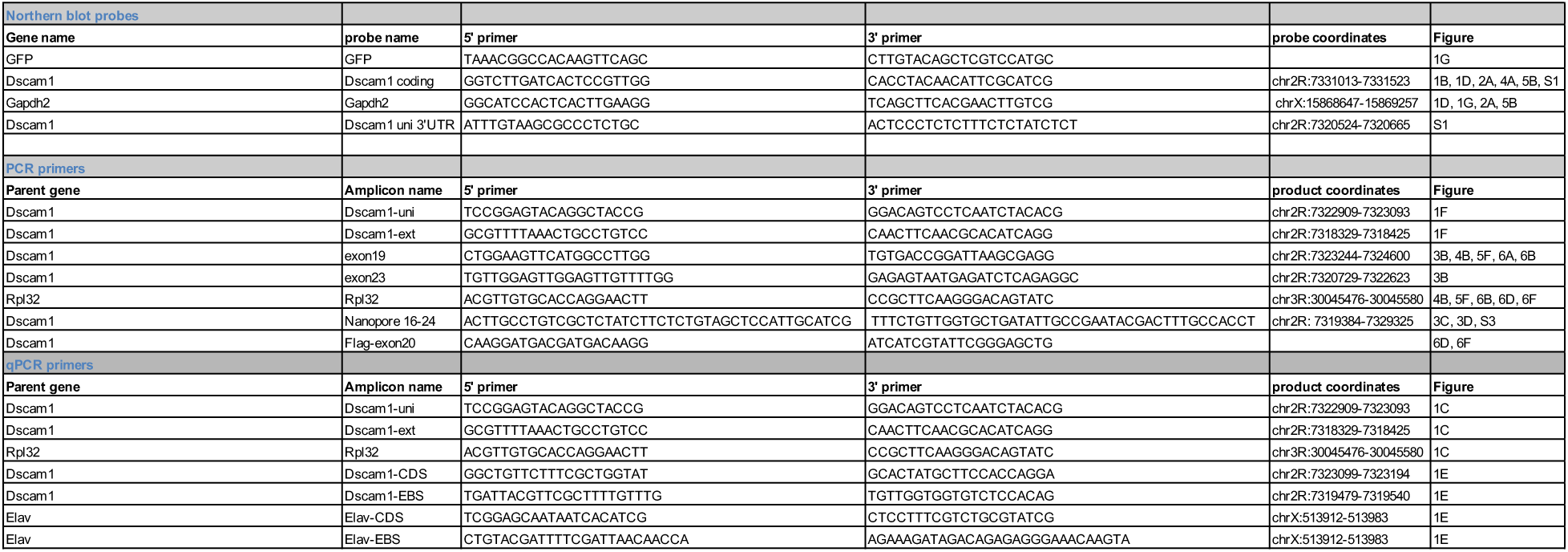
Oligonucleotide sequences. Related to Figures 1, 2, 3, 4, 5, 6. Coordinates correspond to *D. Melanogaster* dm6 genome.

## Supplemental Multimedia Files

**Video #1. Related to** Figures 2 **and** 4. *Elav-GAL4/+; UAS-Dicer2/UAS-shExt* flies display impaired locomotion and could not fly. Flies were placed on pad without anesthesia.

**Video #2. Related to Figure 4.** *Elav-GAL4/+; UAS-Dicer2/UAS-sh-19* flies display impaired locomotion and could not fly. Flies were placed on pad without anesthesia.

## KEY RESOURCES TABLE

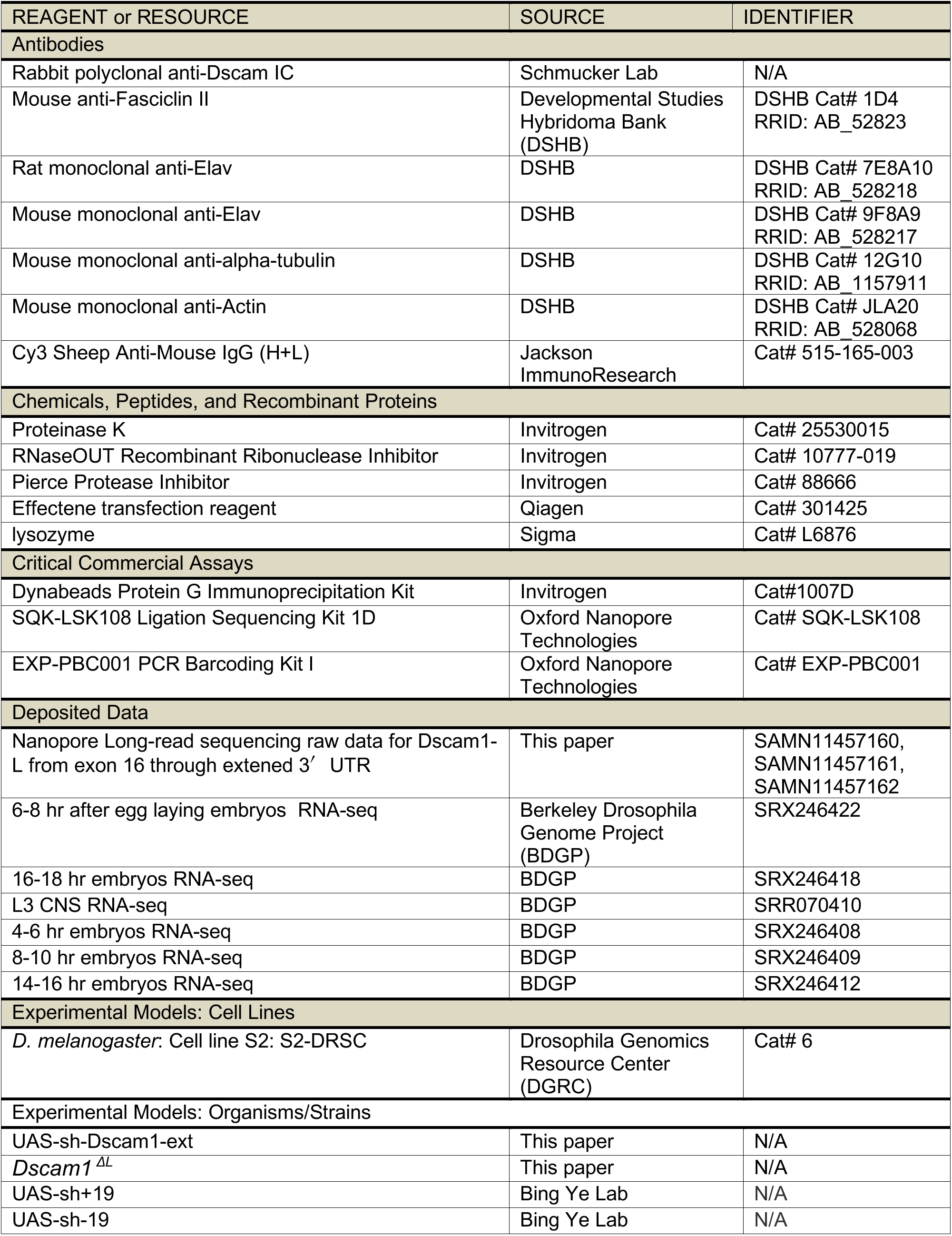
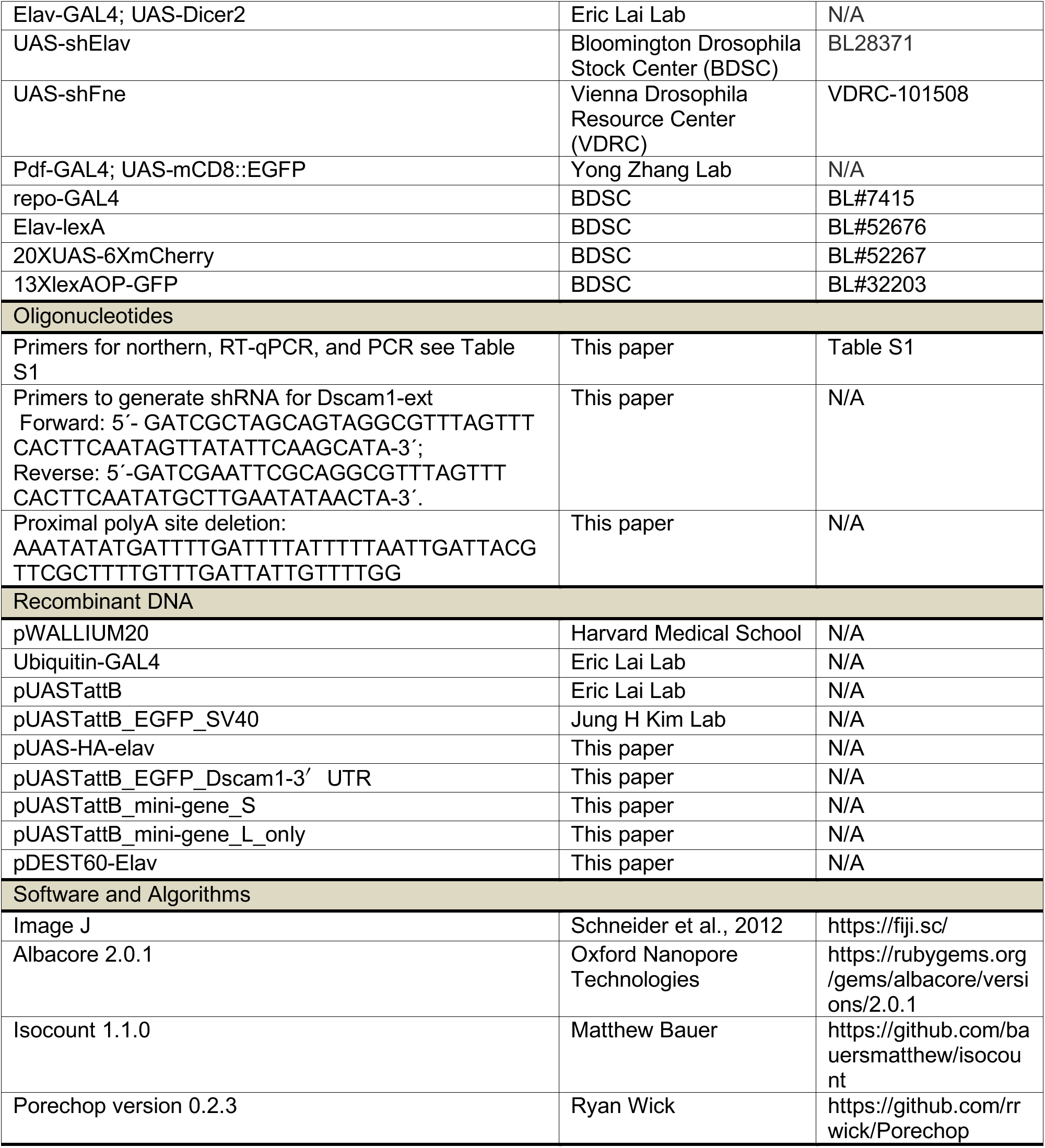

